# The cell adhesion molecule Echinoid promotes tissue survival and separately restricts tissue overgrowth in *Drosophila* imaginal discs

**DOI:** 10.1101/2023.08.04.552072

**Authors:** Danielle C. Spitzer, William Y. Sun, Anthony Rodríguez-Vargas, Iswar K. Hariharan

## Abstract

**The interactions that cells in *Drosophila* imaginal discs have with their neighbors are known to regulate their ability to survive. In a screen of genes encoding cell surface proteins for gene knockdowns that affect the size or shape of mutant clones, we found that clones of cells with reduced levels of *echinoid* (*ed*) are fewer, smaller, and can be eliminated during development. In contrast, discs composed mostly of *ed* mutant tissue are overgrown. We find that *ed* mutant tissue has lower levels of the anti-apoptotic protein Diap1 and has increased levels of apoptosis which is consistent with the observed underrepresentation of *ed* mutant clones and the slow growth of *ed* mutant tissue. The eventual overgrowth of *ed* mutant tissue results not from accelerated growth, but from prolonged growth resulting from a failure to arrest growth at the appropriate final size. Ed has previously been shown to physically interact with multiple Hippo-pathway components and it has been proposed to promote Hippo pathway signaling, to exclude Yorkie (Yki) from the nucleus, and restrain the expression of Yki-target genes. We did not observe changes in Yki localization in *ed* mutant tissue and found decreased levels of expression of several Yorkie-target genes, findings inconsistent with the proposed effect of Ed on Yki. We did, however, observe increased expression of several Yki-target genes in wild-type cells neighboring *ed* mutant cells, which may contribute to elimination of *ed* mutant clones. Thus, *ed* has two distinct functions: an anti-apoptotic function by maintaining Diap1 levels, and a function to arrest growth at the appropriate final size. Both of these are unlikely to be explained by a simple effect on the Hippo pathway.**

## Introduction

Nowhere is the influence of neighbors more important than for cells in developing epithelia. In unicellular organisms, the availability of nutrients is the primary regulator of cell proliferation. In addition to nutrient availability, the survival and proliferation of epithelial cells in multicellular organisms is regulated by a variety of signals emanating from other cells. Such signals include diffusible factors that could be secreted locally or circulate systemically. The packing of cells into tissue layers generate mechanical forces that can influence cell proliferation and survival. Additionally, especially in epithelial tissues, contacts between cells and their immediate neighbors, likely mediated by cellular adhesion molecules, can determine whether cells remain within the layer or get extruded; extrusion often accompanies or promotes cell death.

The importance of a cell’s immediate neighbors in regulating its survival and proliferation is best illustrated by the phenomenon of cell competition (Morata and Ripoll 1975) (reviewed by Amoyel and Bach 2014; Nagata and Igaki 2018; Baker 2020; Cumming and Levayer 2023). Cell competition does not refer simply to the ability of fast-growing cells to outpace slow-growing cells. Rather, it is a situation where cells that are less fit for a variety of reasons are underrepresented because they are eliminated only in the presence of faster-growing cells (i.e., in genetic mosaics). In their original discovery of this phenomenon, Morata and Ripoll found that cells heterozygous for a class of mutations known as *Minute*, that mostly disrupt ribosomal proteins, are eliminated in the presence of wild-type cells. However, as a homogeneous population, they can generate animals of relatively normal size and shape albeit much more slowly. Subsequently, small patches of wild-type cells were shown to be eliminated when they are surrounded by cells having even modest increases in the level of the Myc protein (Moreno and Basler 2004; de la Cova *et al*. 2004). Cells capable of eliminating wild-type cells were dubbed “supercompetitors”. Additionally, cells with mutations in the Hippo pathway that have increased activity of the transcriptional co-activator Yorkie (Tyler *et al*. 2007; Neto-Silva *et al*. 2010), cells with increased Wnt signaling (Vincent *et al*. 2011), cells with increased Jak/Stat signaling (Rodrigues *et al*. 2012) and cells with reduced *crumbs* function (Hafezi *et al*. 2012) were also shown to be supercompetitors. Collectively these findings show that the designation of “winners’’ and “losers’’ is not determined by the genotype of the cell itself but by how it compares itself to its neighbors. How this comparison is achieved is still not well understood since many of the mutations that alter competitive ability affect intracellular proteins and any kind of comparison must occur at the cell surface. Once the comparison has happened, specific isoforms of the cell surface protein Flower are upregulated in “loser” cells (Rhiner *et al*. 2010). Also, winner cells seem capable of expressing higher levels of the Toll ligand Spätzle which might promote the loser cell fate in adjacent cells (Alpar *et al*. 2018).

There are several other phenomena that resemble classical cell competition, some of which appear to be mechanistically distinct. Clones of cells with mutations in genes encoding proteins that regulate apicobasal polarity such as *scribble* (*scrib*) and *discs large* (*dlg*) are eliminated during development (Brumby and Richardson 2003). However imaginal discs or entire compartments of discs that are entirely composed of mutant cells overgrow and the epithelia are often multilayered (Bilder *et al*. 2000). The elimination of clones of *scrib* or *dlg* cells requires the TNF-ortholog Eiger (Igaki *et al*. 2009). The loss of apicobasal polarity re-localizes cell surface proteins including the Eiger receptor, Grindelwald, rendering it accessible to circulating Eiger with the result that cells undergo apoptosis (de Vreede *et al*. 2022). Also, a signaling event at the clone interface involving the ligand Stranded at second (Sas) and the receptor PTP10D that promotes the elimination of polarity-deficient cells has been proposed (Yamamoto *et al*. 2017) but a requirement for PTP10D has been questioned by others (Gerlach *et al*. 2022).

In addition to these phenomena, mis-specified cells can be eliminated by a JNK-dependent pathway (Adachi-Yamada and O’Connor 2002) that involves the transmembrane protein Fish-lips (Adachi-Yamada *et al*. 2005). More recently, there has been an increasing appreciation of the importance of mechanical forces in eliminating cells that proliferate more slowly (Shraiman 2005; Marinari *et al*. 2012; Mao *et al*. 2013; Levayer *et al*. 2016). Such mechanisms might make some cells more susceptible to elimination but this kind of mechanism cannot, on its own, easily account for observations that most, if not all cells, of specific genotypes are selectively eliminated.

One class of proteins that are likely to play an important role in mediating heterotypic interactions at clone interfaces are cell-surface proteins, particularly cell-cell adhesion proteins. The *Drosophila* genome encodes over one hundred proteins containing cadherin motifs or Ig-loops, which are commonly used for cell-cell adhesion (Hynes and Zhao 2000; Vogel *et al*. 2003). Although many of these genes have been studied extensively in neuronal contexts, relatively few have been examined for a role in cell survival or cell proliferation in epithelia (Finegan and Bergstralh 2020), and fewer yet in a mosaic context. Here we describe a genetic screen where we have reduced the function of individual cell-surface proteins in clones of cells in the wing imaginal disc and identified those that affect the size or shape of mutant clones. Of these, we focus on the properties of the Echinoid (Ed) protein because its depletion in clones results in clone elimination while reducing *ed* function in the entire disc results in overgrowth. We demonstrate a role of Ed in maintaining levels of the anti-apoptotic protein Diap1 in cells, and also a separate role in arresting growth when a tissue reaches its final size.

## Results

The overall goal of our screen was to identify cell adhesion molecules that regulated the survival, proliferation, or arrangement of epithelial cells in imaginal discs. To that end we generated clonal patches of cells in wing imaginal discs, that had reduced levels of individual adhesion molecules, in the midst of wild-type cells, and assessed the number, size and shapes of the mutant clones. We used the FLP-out Gal4 system (Pignoni and Zipursky 1997) to activate the expression of Gal4 in random isolated cells and their progeny, and concurrently expressed an RNAi transgene under the control of Gal4-responsive *UAS* elements (Fischer *et al*. 1988; Brand and Perrimon 1993). The cells expressing the RNAi transgene were marked by concurrent expression of a fluorescent protein (*UAS-GFP* or *UAS-RFP*). The size and shape of clones were compared to those expressing an RNAi transgene directed against the *white* (*w*) gene (*UAS-w-RNAi)*.

For our screen, we initially compiled a list of 151 genes that encoded known or putative cell-adhesion molecules, based on the assumption that they had cadherin or immunoglobulin domains (Figure 1A) (Supplementary Table 1). These included those identified by Hynes & Zhao (2000) or Vogel et al. (2003) as well as others that we included based on reports in the literature. Since we chose to conduct our screen in the wing imaginal disc, we excluded 43 genes that were not expressed in wing discs using the single-cell RNAseq data of Everetts et al. (2021) and 9 genes whose localization and function precluded a role in cell-cell adhesion (e.g. sarcomere components) (Supplementary Table 2). Of the remaining 99 genes, we screened 74 genes using 90 different RNAi lines (Supplementary Table 3).

**Figure 1:**
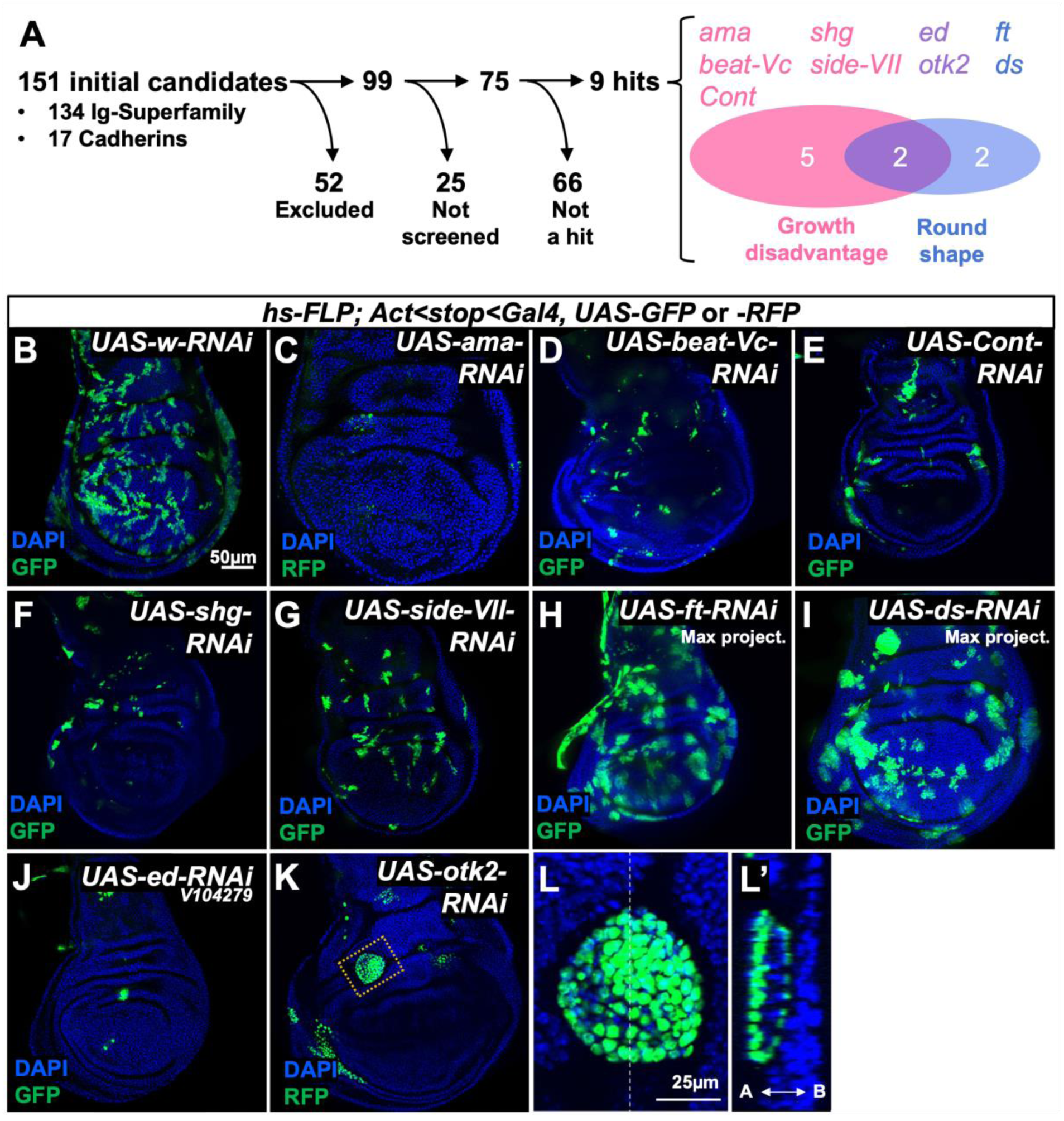
Clonal phenotypes observed in RNAi screen. (**A**) Summary of the screen where genes encoding cell-surface proteins were individually knocked down using RNAi transgenes (**B-L**) Phenotypes of imaginal discs containing clones generated with a FLP-out Gal4 and UAS-RNAi transgenes, Clones are marked by the inclusion of a *UAS-GFP* or *UAS-RFP* transgene as indicated. A cyst-like clone in (**K**) is shown at higher magnification in (**L**) and an orthogonal view in (**L’**). (**B-K**) are shown at the same magnification; the scale bar, 50 μm is shown in panel (**B**). (**L, L’**) are shown at a higher magnification: scale bar is 25 μm.

While expression of most of the RNAi lines did not cause an obvious change in the number, size, or shape of the clones when compared to the expression of *w-RNAi* (Figure 1B), five lines caused a reduction in clone size, two affected clone shape without reducing clone size, and two reduced clone size and also generated rounder clones (Figure 1A). Those that caused a marked reduction in the number and size of clones were *amalgam* (*ama*), *beaten path Vc* (*beat-Vc*), *sidestep VII* (*side-VII*), *Contactin* (*Cont*), and *shotgun* (*shg*) (Figure 1C-G). Amalgam is a secreted protein whose best characterized function is in axonal fasciculation where it promotes the adhesion of cells and axons that express Neurotactin to each other (Frémion *et al*. 2000). Beat and Side family proteins interact with each other to facilitate cell adhesion. A comprehensive analysis of physical interactions between Beat and Side proteins did not predict an interaction between Beat-Vc and Side-VII; Side-VII was predicted to bind to Beat-IV and Beat-Vc was predicted to bind to Side-VI (Li *et al*. 2017). Contactin is a GPI-anchored protein needed to organize septate junctions of epithelial cells (Faivre-Sarrailh *et al*. 2004). *shg* encodes the *Drosophila* ortholog of E-cadherin that functions as a homophilic adhesion molecule that is a key component of adherens junctions (Oda *et al*. 1994; Tepass *et al*. 1996). Clones expressing RNAi constructs targeting *fat* (*ft*) (Figure 1H) and *dachsous* (*ds*) (Figure 1I) are clearly rounder and likely larger than *w-RNAi* clones. Fat and Dachsous are large atypical cadherins whose extracellular domains bind to each other and their role in regulating cell proliferation and the orientation of cell division has been studied extensively (reviewed by Fulford & McNeill, 2020; Thomas & Strutt, 2012). Knockdown of two genes, *echinoid* (*ed*) (Figure 1J) and *off-track2* (*otk2*) (Figure 1K, L) generated fewer clones that were typically smaller and also appeared more rounded when compared to the irregularly-shaped elongated clones that are normally found in the wing pouch (Figure 1B). Otk2 clones were sometimes extruded as cysts (Figure 1L, L’). The function of Otk2 is not well understood other than it likely functions as a co-receptor for the Wnt2 ligand (Linnemannstöns *et al*. 2014).

Clones of cells with reduced *ed* function were infrequent, small, and had smooth outlines. The reduced growth of *ed* clones in imaginal discs has been noted previously (Escudero *et al*. 2003; Wei *et al*. 2005) but is not easily reconciled with some of the other functions of Ed (see below). We therefore chose to characterize the function of Echinoid in growth regulation in more detail. The Echinoid protein has an extracellular domain that includes seven Ig repeats, three FN-III repeats and an intracellular domain of 315 amino acids that includes a C-terminal PDZ-binding protein that can interact either with Bazooka (Baz) or Canoe (Canoe) (Bai *et al*. 2001; Wei *et al*. 2005). *ed* mutant clones are rounded, in contrast to the irregular (“wiggly”) boundaries of wild-type clones (Wei *et al*. 2005). *ed* mutant cells fail to assemble proper adherens junctions (AJs) at the interface wild-type cells (Wei *et al*. 2005; Laplante and Nilson 2006; Chang *et al*. 2011). This differential adhesiveness might result in the tendency for these cells to remain together and sort away from wild-type cells (Townes and Holtfreter 1955; Steinberg 1963). Additionally, an actomyosin cable that forms at Ed expression boundaries in the wild-type cells is likely to act as a “mechanical fence” at the clone perimeter (Wei *et al*. 2005; Laplante and Nilson 2006, 2011; Lin *et al*. 2007; Chang *et al*. 2011). Together, these properties explain the roundness and smoothness of *ed* clones at apical levels.

Ed is also thought to function as a signaling molecule. *ed* mutations were originally identified as dominant enhancers of a hypermorphic allele of the EGF receptor, *Ellipse* (*Elp*) (Bai *et al*. 2001). Ed has subsequently been shown to be a negative regulator of EGF receptor (EGFR) signaling (Bai *et al*. 2001; Islam *et al*. 2003; Rawlins *et al*. 2003a; Spencer and Cagan 2003; Fetting *et al*. 2009; Ho *et al*. 2010) and a positive regulator of Notch signaling (Ahmed *et al*. 2003; Escudero *et al*. 2003; Rawlins *et al*. 2003b), possibly by promoting endocytosis of EGFR and Delta respectively. More recently, reduced *ed* function has also been shown to cause tissue overgrowth by decreasing signaling via the Hippo pathway and increasing the levels of nuclear Yorkie (Yki) (Yue *et al*. 2012). Ed interacts with multiple Hippo -pathway components, promotes the stability of Salvador (Sav) and curtails the expression of Yki-target genes that normally promote growth and cell survival. Since both the EGFR pathway and Yki promote cell survival and cell proliferation (Díaz-Benjumea and García-Bellido 1990; Bergmann *et al*. 1998; Kurada and White 1998; Huang *et al*. 2005), and both EGFR signaling and Yki-target gene expression would be expected to be elevated in *ed* mutant tissue, the underrepresentation of *ed* mutant tissue is not easily explained. We therefore decided to examine the properties of *ed* mutant tissue in greater detail.

### Clones of *echinoid* mutant cells in epithelia are eliminated by a process that resembles cell competition

To avoid the possibility of off-target effects generated with the *ed-RNAi* transgene used in our screen (V104279), we also tested three other *ed-RNAi* lines. In each case, clones were induced 72 h after egg lay (AEL) and imaginal discs were dissected 48 h later at 120 h AEL. Both V104279 (Figure 2A) and V3087 (Figure 2B) generated fewer and smaller clones and Ed protein was almost undetectable in those clones. The BL38243 line generated clones that were not smaller and had wiggly outlines, much like wild-type clones (Figure 2D); the knockdown was least effective in this line (Figure 2D’, 2D’’). The V938 line had an intermediate effect (Figure 2C, C’, C’’). We also used mitotic recombination with the MARCM method where mutant clones are positively marked with GFP (Figure 2E-H). Using this approach, homozygous mutant clones of two different null alleles of *ed* (*ed^IF20^* and *ed^1×5^*) (de Belle *et al*. 1993; Bai *et al*. 2001), that have early stop codons and would be predicted to encode proteins with small portions of the extracellular domain (Escudero *et al*. 2003), were completely absent from the disc epithelium (Figure 2F-G). Clones that are homozygous for the hypomorphic allele, *ed^sIH8^* (de Belle *et al*. 1993), which has a missense mutations that changes a conserved cysteine residue in the sixth Ig domain of the extracellular portion to serine (Escudero *et al*. 2003), are observed but are also far less frequent and smaller than wild-type clones (Figure 2H). Notably, in all of these experiments, large clones of myoblasts are often observed beneath the epithelium in the notum region of the disc (Figure 2F’). Although Ed is expressed in wing disc-associated myoblasts based on single cell RNA-seq data (Everetts *et al*. 2021) and is required for the targeting and morphogenesis of some embryonic and larval body wall muscles (Swan *et al*. 2006), unlike *ed*-depleted epithelial cells, *ed*-depleted myoblasts do not have a growth or survival disadvantage.

**Figure 2:**
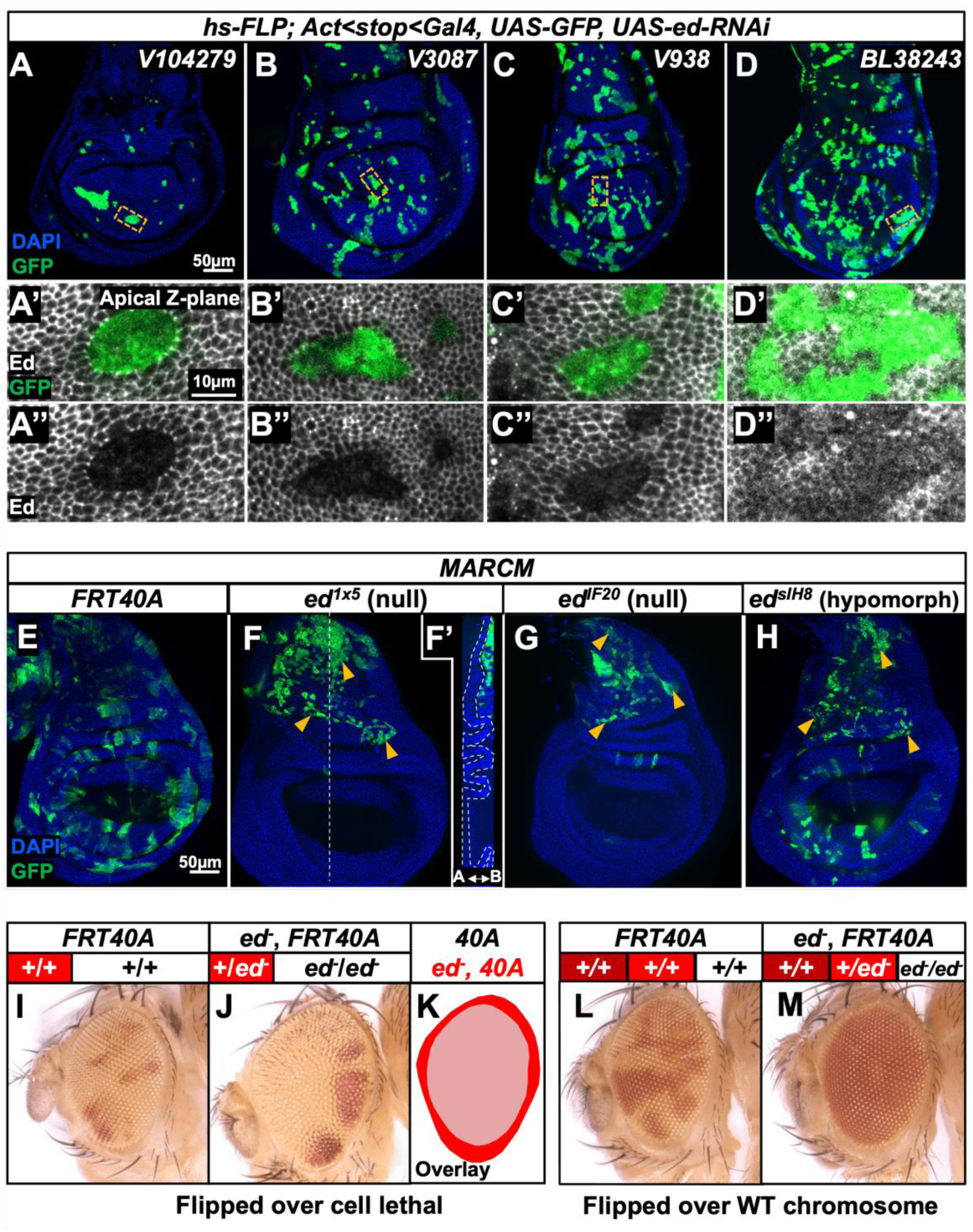
Clones of *echinoid* mutant cells are eliminated when surrounded by wild-type cells. (**A-D**) Imaginal discs containing GFP-marked clones of cells that express an RNAi directed against *ed*. Four different UAS-RNAi transgenes were used. Panels (**A’-D’**) and (**A’’-D’’**) show GFP-marked clones at higher magnification that are stained with an anti-Ed antibody. The magnified regions are indicated with dashed lines in (**A-D**). (**E-H**) GFP-marked clones of cells that are homozygous for the chromosome arm bearing *FRT40A* alone (**E**) or *FRT40A* and an allele of *ed* (**F-H**). Note the near absence of clones in the epithelium with the two null alleles of *ed* (**F, G**) and the presence of myoblast clones underlying the notum (indicated by an arrowheads). An orthogonal view is shown in (**F’**) with the disc epithelium outlined; the GFP-marked myoblasts are located basal to the epithelium. Both epithelial and myoblast clones are observed with the hypomorphic allele (**H**). (**I-K**) Clones of cells homozygous for either a wild-type chromosome arm distal to *FTR40A* (**I**) or *ed^IF20^, FRT40A* (**J**) marked white, generated using *eyFLP*. The tester chromosome carries a recessive cell lethal resulting in the absence of wild-type twin spots. An overlay of the overgrown eye containing *ed* clones and the normally sized eye containing wild-type clones is shown in (**K**). (**L, M**) Clones generated using *eyFLP* using a wild-type tester chromosome that does not carry a recessive cell lethal mutation. *FRT40A* clones (**L**) or *ed FRT40A* clones (**M**) are white while the wild-type twin spots appear red in both panels. Note the almost complete absence of homozygous *ed/ed* tissue in eyes that contain wild-type twin spots.

Yue et al. (2012) had reported that *ed/ed* clones generated during eye development were overgrown. An important difference between that experiment and our experiments is that their clones were generated in a stock where the wild-type chromosome carried a recessive cell-lethal mutation. Thus, following *eyFLP*-driven mitotic recombination, the eye disc would contain mostly *ed/ed* tissue as well as a small amount of *ed/+* tissue that had not undergone mitotic recombination. Consistent with their observations, we find that eyes composed almost entirely of *ed^IF20^/ed^IF20^*tissue are overgrown (Figure 2I-K). When the wild-type twin clones are not eliminated with a recessive cell-lethal, we observed almost no homozygous *ed/ed* tissue (Figure 2L-M). Thus *ed/ed* tissue is able to survive, and even overgrow, when the disc is mostly composed of *ed/ed* tissue. In contrast, *ed/ed* tissue is extremely underrepresented in the presence of wild-type tissue.

To further explore this phenomenon, we generated discs where an increasing proportion of tissue had decreased *ed* function by increasing the density of clones expressing *ed-RNAi*. This was achieved by expressing *hs-FLP* for 12 min, 15 min, or 30 min (Figure 3A-F). At low clone densities, *ed-RNAi*-expressing clones were far fewer than *w-RNAi* (control) clones (Figure 3A, B). As clone density increased, until the clones collectively accounted for the majority of tissue in the wing disc, the representation of *ed-RNAi* tissue and *w-RNAi* tissue was similar (Figure 3C-F). Thus, the loss of *ed* mutant tissue is most likely to occur when those clones are completely surrounded by wild-type cells.

**Figure 3:**
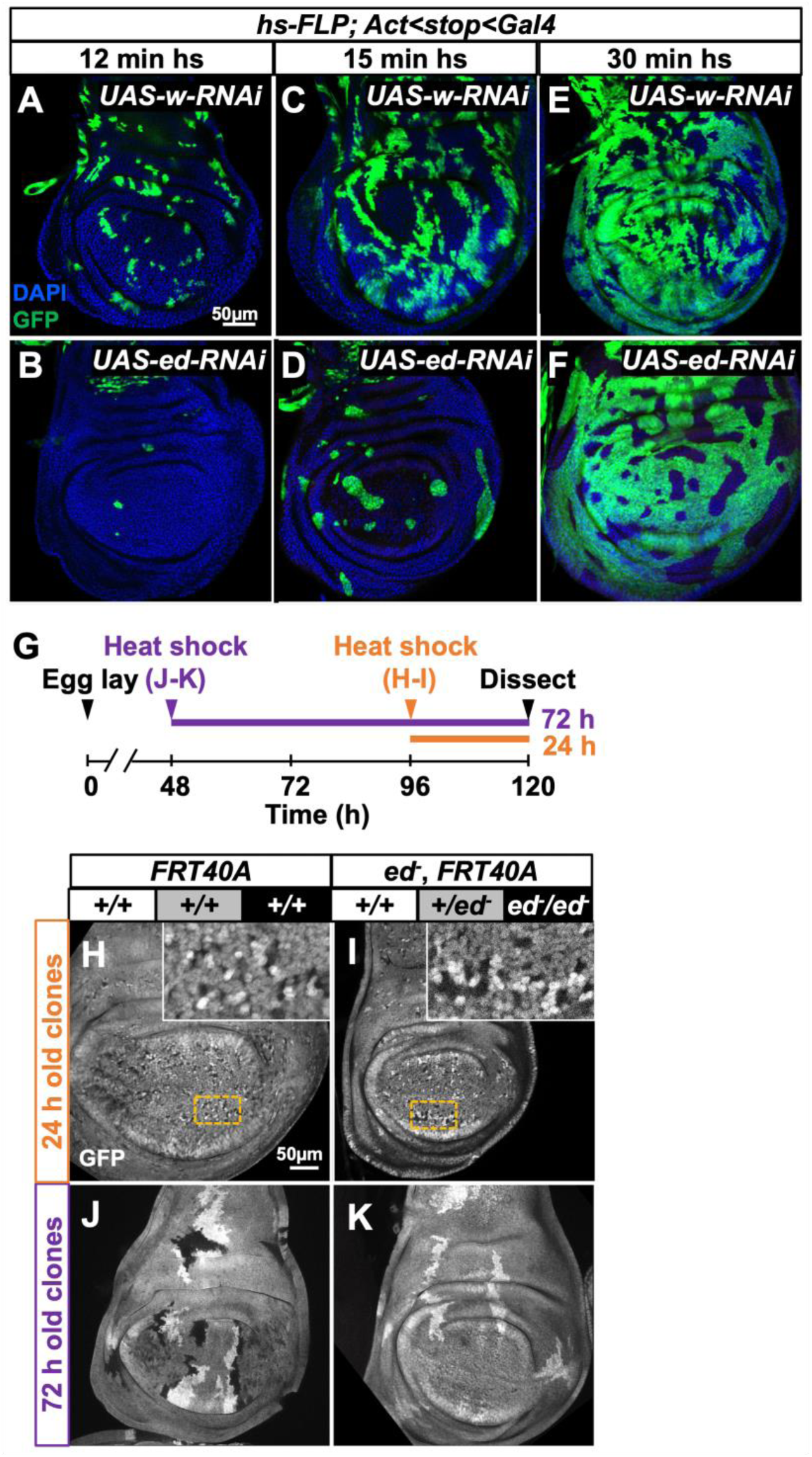
*echinoid* clones are generated and then die, especially when more wild-type tissue is present. (**A-F**) Wing discs containing clones expressing either *UAS-w-RNAi* (**A, C, E**) or *UAS-ed-RNAi* (**B, D, F**). Clones also express GFP. The clone density was varied by increasing the duration of the heat shock at 37° C during which *hs-FLP* is expressed. Thus, heat shocks of 12 min (**A, B**), 15 min (**C, D**) and 30 min (**E, F**) generate clones at progressively higher density. (**G-K**) An experiment to examine *ed/ed* clones 24 h and 72 h after generation by mitotic recombination. The design of the experiment is shown in (**G**). Discs were dissected 120 h AED and clones were induced either 24 h (**H, I**) or 72 h (**J, K**) prior to dissection. Clones generated using the *FRT40A* chromosome (**H, J**) are compared to clones homozygous for *ed^IF20^* (**I, K**).

In their original description of cell competition, Morata & Ripoll (1975) observed that *Minute* clones were eliminated when they were generated early in development but were still observable when they were generated much later. To determine if this was also the case with clones of *ed/ed* tissue, we generated clones either 48 h or 96 h AEL using mitotic recombination and discs were dissected at 120 h AEL (Figure 3G). Thus, the discs were allowed to develop for 72 h or 24 h after clone induction respectively (Figure 3H-K). With marked wild-type clones (Figure 3H, J), clones were much larger 72 after clone generation (Figure 3J) than they were 24 h after clone generation (Figure 3H). In marked contrast *ed/ed* clones were observed readily 24 h after clone generation (Figure 3I) but mostly absent 72 h after clone generation (Figure 3K). At 72 h after clone generation their wild-type sister clones were similar in size to wild-type clones generated at similar times (Figure 3J-K). Thus *ed/ed* clones proliferate for a short time and are then eliminated. The propensity of *ed/ed* tissue to be eliminated when surrounded by wild-type tissue and to survive and grow when it accounts for most of the tissue is very similar to other instances of cell competition.

### Increased apoptosis and reduced levels of Diap1 are observed in *echinoid* mutant tissue

In many situations where clones of cells of a certain genotype (e.g. *Minute* heterozygotes) are eliminated by cell competition, their elimination occurs by caspase-mediated apoptotic cell death and their death can be rescued by expression of the baculovirus p35 protein (Martín *et al*. 2009) which is an inhibitor of effector caspases (Hay *et al*. 1994). We therefore generated clones expressing either *w-RNAi* or *ed-RNAi* either in the presence or absence of *p35* (Figure 4A-D). While expression of *p35* modestly increases the recovery of wild-type clones (Figure 4A, B), there was a dramatic increase in the recovery of clones expressing *ed-RNAi* (Figure 4C, D).

**Figure 4:**
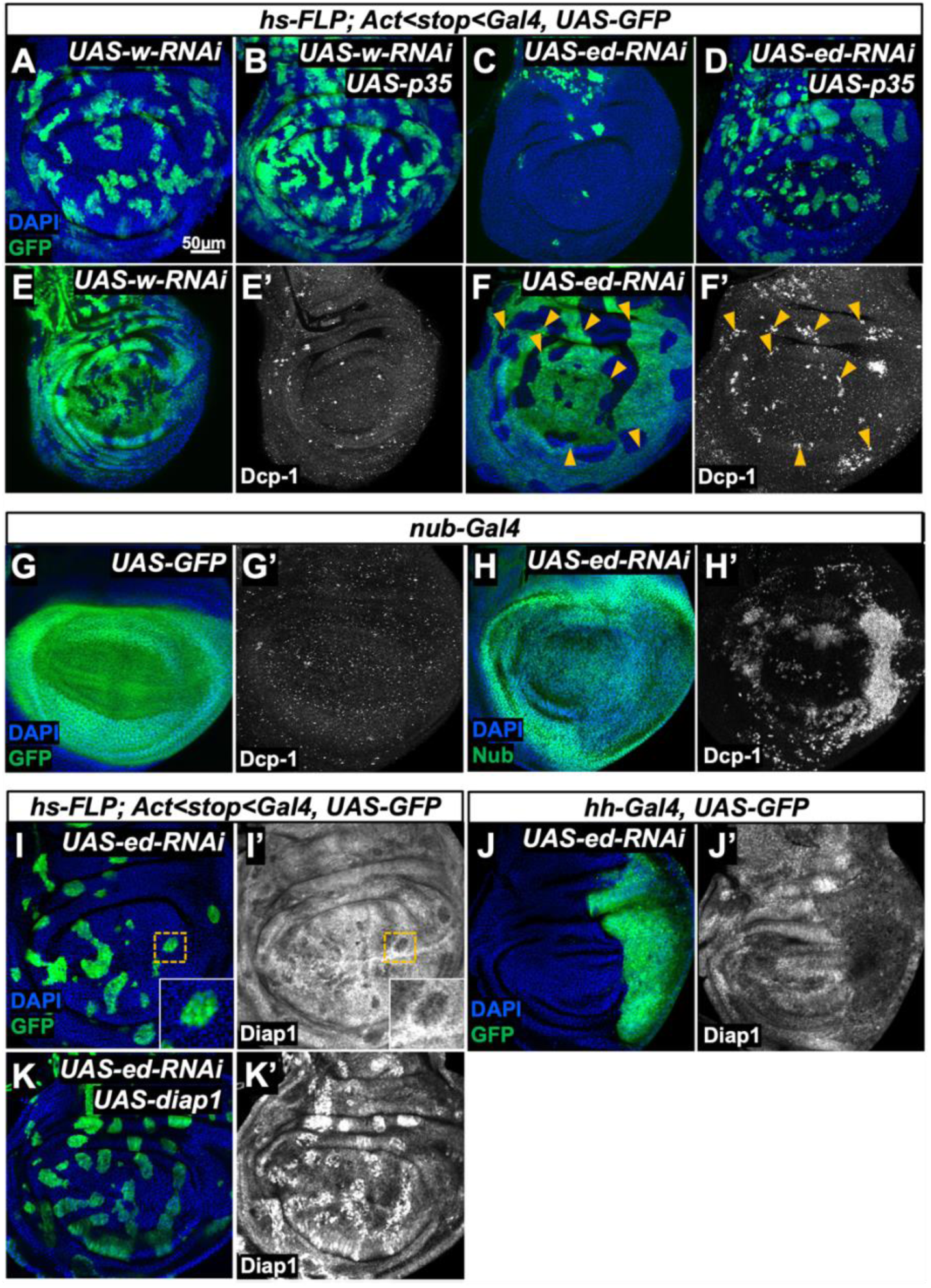
*echinoid* mutant tissue has lower Diap1 levels and has higher levels of apoptosis. (**A-D**) Wing discs containing GFP-marked clones of cells that express either *UAS-w-RNAi* (**A, B**) or *UAS-ed-RNAi* (**C, D**). Clones that also express *UAS-p35* are shown in (**B, D**). (**E, F**) Imaginal discs containing GFP-marked clones expressing *UAS-w-RNAi* (**E**) or *UAS-ed-RNAi* (**F**). Anti Dcp-1 antibody staining of the same discs is shown in (**E’, F’**). Images were taken at a basal Z-plane where Dcp-1 staining was most prominent. (**G, H**) Effect of reducing *ed* function in the entire wing pouch on levels of apoptosis. *nub-Gal4* drives expression of *UAS-GFP* in (**G, G’**) or *UAS-ed-RNAi* in (**H, H’**). The pouch is visualized by GFP expression in (**G**) and with anti-Nubbin in (H). Apoptotic cells are visualized with anti-Dcp1. Note increased levels of apoptosis at the periphery of the pouch where **nb-Gal4** expression is higher. (**I, J**) Discs expressing *ed-RNAi* in clones (**I**) or in the entire posterior compartment (**J**) stained with an anti-Diap1. The insets in (**I, I’**) show a clone at higher magnification. (**K**) Discs with GFP-marked clones that express both *UAS-ed-RNAi* and *UAS-diap1* which results in an increase in both the number and size of clones. Staining with anti-Diap1 is shown in (**K’**). Note that the “rescued” clones still have smooth borders consistent with the propensity of cells with decreased *ed* to sort away from wild-type cells.

These observations indicate that the activity of effector caspases is necessary for the elimination of *ed* mutant tissue. Apoptotic cells in *Drosophila* express Death Caspase-1 (Dcp-1) (Song *et al*. 1997). To visualize cells undergoing apoptosis, we used the anti-Dcp-1 antibody. In discs containing clones expressing *w-RNAi*, occasional Dcp-1 staining was observed (Figure 4E, E’). In contrast, in discs containing *ed-RNAi* clones, punctate Dcp-1 staining was observed within *ed-RNAi* clones, especially in basal focal planes (Figure 4F, F’). These puncta were often at the edge of the clone where it abuts wild-type tissue, suggesting that competition with neighbors may be involved. We also examined Dcp-1 staining when using the *nub-Gal4* driver. Very little apoptosis was seen when *nub*-*Gal4* drove the expression of GFP (Figure 4G, G’), but very high levels of apoptosis were seen in the pouch of *nub-Gal4, UAS-ed-RNAi* discs (Figure 4H, H’). Taken together, these observations indicate that cells with reduced *ed* function die by apoptosis at an increased rate, and this contributes, in significant part, to the elimination of *ed* clones.

An important regulator of apoptosis is the IAP protein Diap1 (Hay *et al*. 1995) which inhibits caspase activation. When discs containing *ed-RNAi* clones were stained with an anti-Diap1 antibody, a clear reduction in Diap1 levels was observed within the clones (Figure 4I, I’). Intriguingly, the wild-type cells immediately adjacent to the clones had elevated levels of Diap1. The elevation of Diap1 in neighboring wild-type cells was inconsistent in that it was not always observed throughout the entire border of the clone, or around every clone. Importantly, a reduction in Diap1 levels was observed not just in *ed* mutant clones, but also when *ed* function was decreased in an entire region of the disc such as the posterior compartment (Figure 4J, J’).

If the reduced level of Diap1 had a role in promoting the death of *ed* mutant cells, then restoring Diap1 levels should be able to prevent clone elimination. To examine this possibility, we expressed a *UAS-diap1* transgene concurrently with *UAS-ed-RNAi*. This resulted in the recovery of many clones that expressed *UAS-ed-RNAi*, an effect that was comparable to that obtained by expressing *p35* (Figure 4K, K’). This indicated that the reduced Diap1 protein level in *ed* mutant clones has a role in their elimination.

The finding that Diap1 levels were consistently reduced in *ed* clones was surprising because a previous study reported that there is decreased activity of the Hippo signaling pathway and increased expression of Yorkie target genes including *diap1* (Yue *et al*. 2012). Clones mutant for components of the Hippo pathway have increased levels of Diap1 protein (Tapon *et al*. 2002) and increased expression of transcriptional reporters of *diap1* (Wu *et al*. 2003). Our finding of decreased Diap1 protein levels in *ed* mutant tissue is inconsistent with the proposed role of Ed promoting Hippo pathway activity; there needs to be another mechanism by which Ed positively regulates Diap1 protein levels. Importantly, our observation of decreased Diap1 in *ed* mutant clones is consistent with their elimination by apoptosis.

### Imaginal discs with reduced *echinoid* function grow more slowly but fail to arrest their growth at the appropriate size

Previous work has indicated that reducing *ed* function in most, or all, cells of the imaginal disc results in overgrowth (Bai *et al*. 2001; Yue *et al*. 2012). This, at first glance, seems inconsistent with the increased levels of cell death observed in *ed* mutant tissue. To analyze the phenomenon further in the wing disc, we first examined the effect of reducing *ed* function on the size of adult wings (Figure 5A-I). Adult flies that were homozygous for the null alleles of *ed* were not obtained. We therefore examined the size of adult wings in a heteroallelic combination, *ed^IF20^*/*ed^sIH8^*. *ed^IF20^* is a null allele, while *ed^sIH8^* is a hypomorphic allele. Wings of these flies were 9% larger than the wings of wild-type (*Oregon-R*) flies. We also reduced *ed* function in the wing pouch using the *nub-Gal4* driver line and found that this increased wing area by 15% compared to flies with *nub-Gal4* driving *w-RNAi*. Because the *nub-Gal4* chromosome itself appears to reduce wing area (Supplementary Figure 1), all wing-area experiments using *nub-Gal4* were compared to controls that also contained *nub-Gal4*, rather than to *Oregon-R*. Thus, as described previously (Bai *et al*. 2001; Yue *et al*. 2012), reducing *ed* function results in tissue overgrowth.

**Figure 5:**
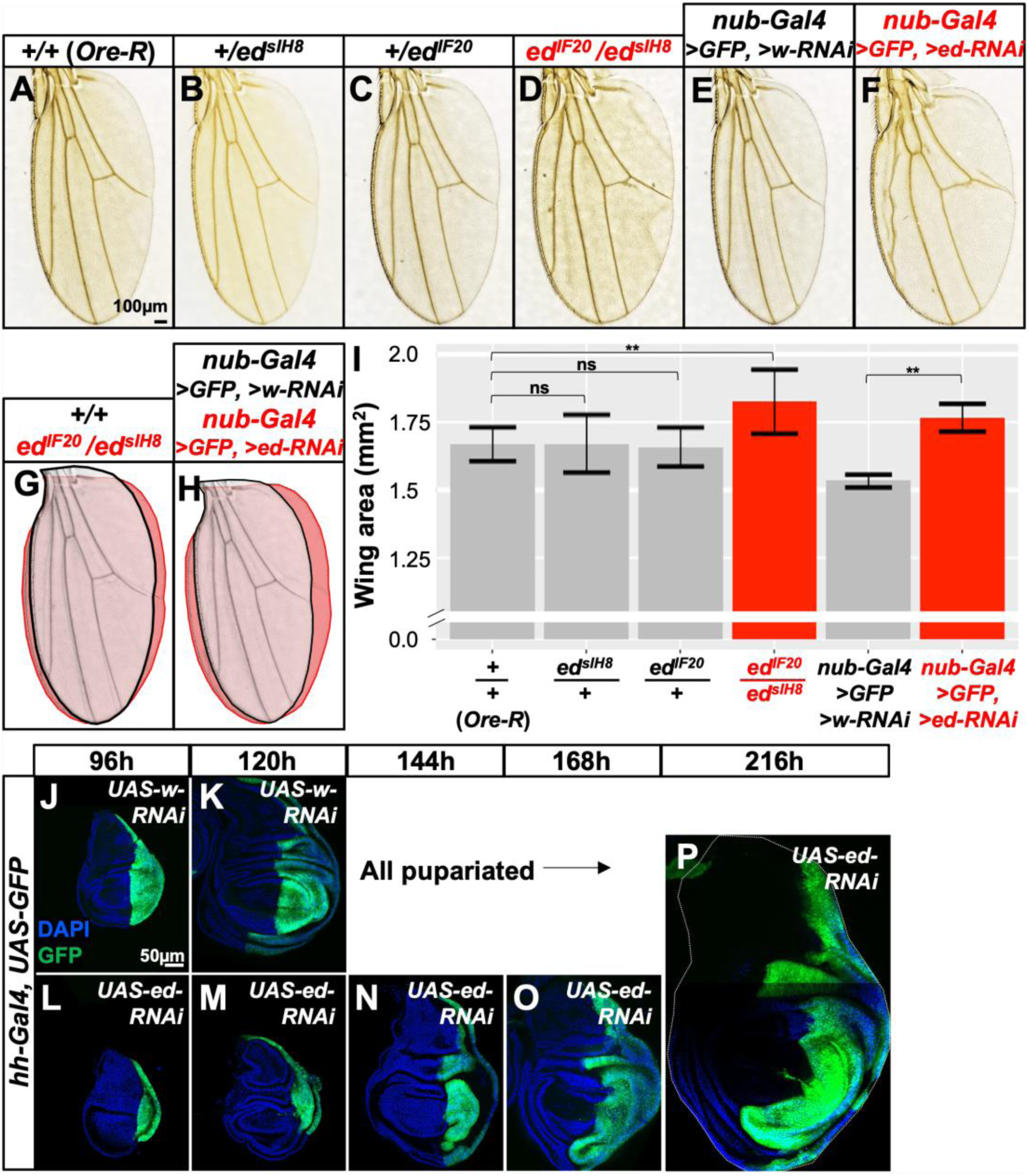
Characteristics of overgrowth exhibited by *echinoid* mutant tissue. (**A-I**) Effect of reducing *ed* function on size of adult wings. (**A-F**) show adult wings of the indicated genotypes. An overlay of a wild-type wing and an overgrown wing of a heteroallelic combination of *ed* mutations is shown in (**G**). An overlay of *nub-Gal4, UAS-w-RNAi* and *nub-Gal4, UAS-ed-RNAi* is shown in (**H**). Quantification of wing areas is shown in (**I**). n=10 wings (*+/+* [*Oregon-R*]*; ed^sIH8^/+; ed^IF20^/+; ed^sIH8^/ed^IF20^; nub-Gal4, >GFP, >w-RNAi*), 9 wings (*nub-Gal4, >GFP, >ed-RNAi*). “ns” indicates p> 0.05, ** indicates p< 0.01, calculated using ANOVA with post-hoc Tukey’s HSD test. Error bars indicate standard deviation. (**J-P**) A time course of growth of imaginal discs expressing either *w-RNAi* (**J, K**) or *ed-RNAi* (**L-P**) in the posterior compartment. All larvae expressing *w-RNAi* have pupariated soon after 120 h. However, much older larvae are observed in the population expressing *ed-RNAi*. Examples of discs from larvae that have delayed their pupariation are shown. All images are shown at the same scale (scale bar in panel **J**).

We then examined the growth of wing discs at different time points during development using the *hh-Gal4* driver which is expressed in the posterior compartment (Figure 5J-P). At 120 h AEL, when control larvae (*hh-Gal4*, *UAS-w-RNAi*) reached the late third instar, the posterior compartments of discs of mutant larvae (*hh-Gal4*, *UAS-ed-RNAi*) were much smaller (Figure 5K, M). However, the discs of mutant larvae continued to grow through an extended larval stage and could reach a size much larger than ever observed in wild-type discs (Figure 5P). Interestingly, although the *ed-RNAi* is only expressed in the posterior compartment, increased growth was also observed in the anterior compartment, possibly as a secondary effect. From these experiments we conclude that, at least under the conditions of this experiment, *ed* mutant tissue grows more slowly than wild-type tissue, likely due to the increased cell death. However, *ed* mutant tissue can continue to grow beyond the appropriate size suggesting a defect in the mechanism that normally arrests growth.

### Mechanisms that function downstream of Echinoid that regulate growth arrest and clone survival

Ed could function in cell-cell adhesion and also as a signaling molecule. To determine which of these properties of Ed are necessary for growth arrest, we examined the ability of either full length Ed protein (encoded by *UAS-ed^Full^*) or one that contains the extracellular domain and the transmembrane domain but lacks the intracellular domain (encoded by *UAS-ed^ΔC-GFP^*) (Laplante and Nilson 2011) to rescue the overgrowth phenotype (Figure 6A-D). We used the heteroallelic combination *ed^sIH8^/ed^IF20^*, since flies of this genotype were viable and, as shown previously (Figure 5A-D, I), their wings were larger when compared to *ed^sIH8^/+* or *ed^IF20^/+* flies. Inclusion of *nub-Gal4* itself resulted in a reduction of wing sizes back to a size similar to that of the wild-type and heterozygous flies (Figure 6A, D; Supplemental Figure 1). Wing size was further reduced by including the *UAS-ed^Full^*transgene and even more so by including the *UAS-ed^ΔC-GFP^*. That *nub-Gal4* on its own caused a reduction in wing size which was exacerbated by either transgene makes it difficult to know whether this is a true rescue of the overgrowth phenotype or whether it is due to a non-specific growth inhibitory effect that counteracts the overgrowth.

**Figure 6:**
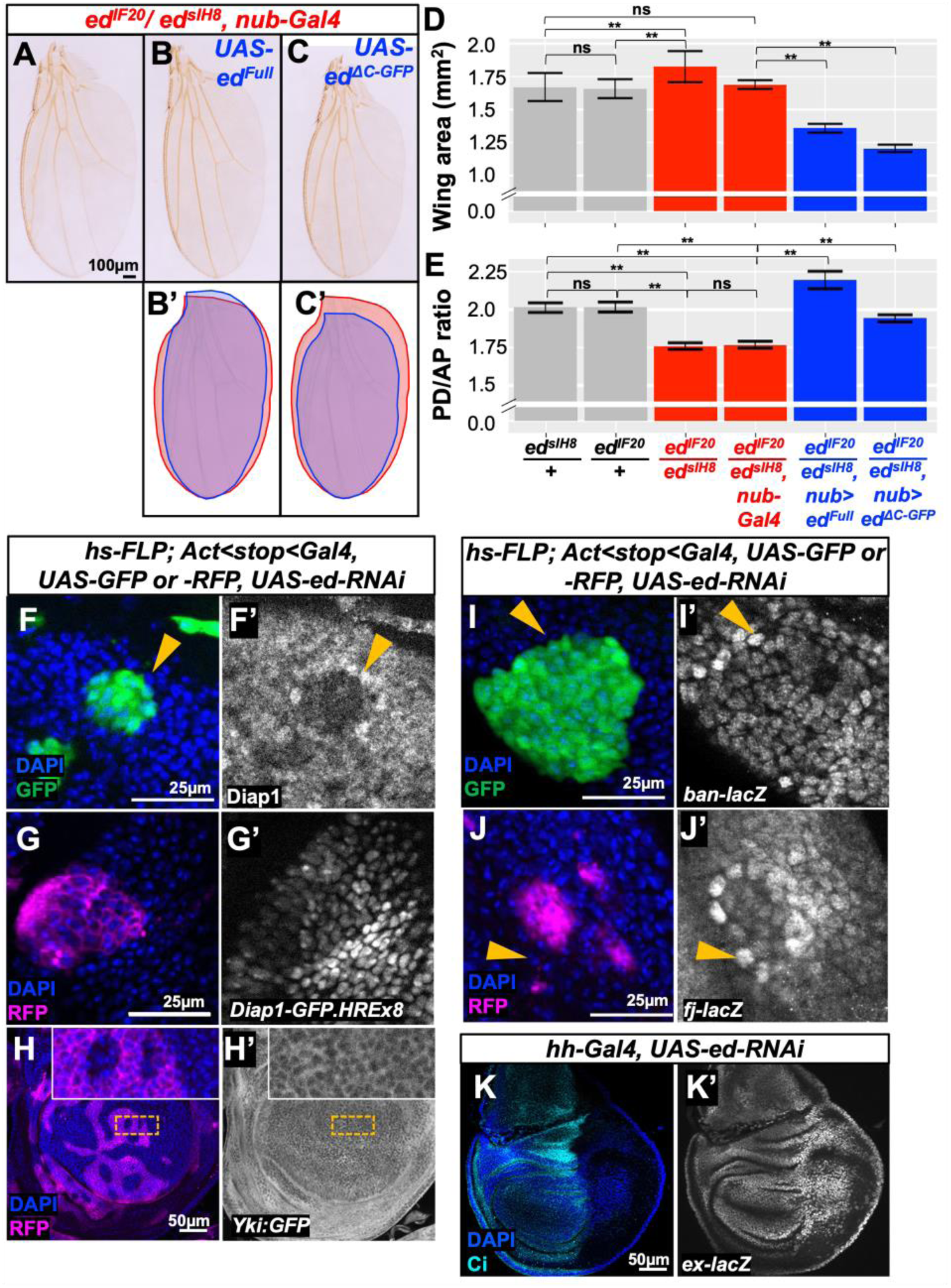
Characterizations of mechanisms that function downstream of *echinoid*. (**A-E**) Effect of expressing full length *ed* (*ed^Full^*) and a version where the cytoplasmic domain has been replaced by GFP (*ed^ΔC-GFP^*) on wing size and wing shape. (**A**) Wings of a heteroallelic combination of *ed* alleles that generate viable adults together with *nub-Gal4*. Inclusion of *UAS-ed^Full^* (**B**) and *UAS-ed^ΔC-GFP^*(**C**) reduces wing size and brings the aspect ratio closer to wild-type. (**B’**) and (**C’**) show overlays of (**B**) and (**C**) over (**A**) respectively. For (**D**), n=10 wings (*ed^sIH8^/+; ed^IF20^/+; ed^sIH8^/ed^IF20^;* same as shown in Fig 5I), 4 wings (*ed^sIH8^/ed^IF20^, nub-Gal4; ed^sIH8^/ed^IF20^, nub-Gal4, UAS-ed^Full^*), 6 wings *(ed^sIH8^/ed^IF20^, nub-Gal4, UAS-ed^ΔC-GFP^*). For (**E**), n=10 wings (*ed^sIH8^/+; ed^IF20^/+*), 9 wings (*ed^sIH8^/ed^IF20^, ed^sIH8^/ed^IF20^, nub-Gal4, UAS-ed^ΔC-GFP^*), 4 wings (*ed^sIH8^/ed^IF20^, nub-Gal4; ed^sIH8^/ed^IF20^, nub-Gal4, UAS-ed^Full^*). The same wings were used in (**D**) and (**E**); number of wings differ if damage or mounting artifacts prevented measurement of both wing area and aspect ratio. “ns” indicates p> 0.05, ** indicates p< 0.01, calculated using ANOVA with post-hoc Tukey’s HSD test. Error bars indicate standard deviation. (**F, F’**) GFP-marked clones expressing *ed-RNAi* stained with anti-Diap1 (**F’**). Yellow arrowhead indicates wild type cells immediately adjacent to the clone. (**G, G’**) RFP marked clone expressing *ed-RNAi* shows decreased expression of a *diap1* transcriptional reporter generated using 8 copies of the Hippo-response element (HRE) from the *diap1* promoter (**G’**). (**H, H’**) RFP-labeled clones expressing *ed-RNAi* show no obvious alteration in the localization of a GFP-tagged Yki. The region of the disc enclosed by the dashed line is shown at higher magnification in the insets. (**I, I’**) GFP-labeled clone expressing *ed-RNAi* shows decreased expression of a *ban-lacZ* reporter (**I’**). The yellow arrowhead indicates wild-type cells adjacent to the clone that show increased expression of *ban-lacZ*. (**J, J’**) RFP-labeled clone expressing *ed-RNAi* shows decreased expression of a *fj-lacZ* reporter (**J’**). The yellow arrowhead indicates wild-type cells adjacent to the clone that show increased expression of *ban-lacZ*. (**K, K’**) Expression of *ed-RNAi* in the posterior compartment of the wing disc results in increased *ex-lacZ* expression (**K’**). The posterior compartment is identifiable because it does not express Ci (**K**).

Flies with *nub-*driven *ed-RNAi* have rounder wings due to a hexagonal packing defect during pupal wing elongation (Chan *et al*. 2021). We also observed rounder wings in *ed^sIH8^/ed^IF20^* flies (Figure 6E). While addition of *nub-Gal4* alone did not change the aspect ratio of *ed^sIH8^/ed^IF20^*flies, *nub-Gal4* driving expression of *UAS-ed^Full^* reduced the roundness of *ed^sIH8^/ed^IF20^* wings. *nub-Gal4* driving expression of *UAS-ed^ΔC-GFP^* gave a significant, though less pronounced, reduction in roundness. This result suggests that Ed is functioning in cell-cell adhesion in regulating organ size and shape. However, since the hypomorphic allele *ed^sIH8^* encodes a protein with a disruption in the extracellular domain, we cannot exclude the possibility that this protein might form a heterodimer with the Ed*^ΔC-GFP^* protein, and that this heterodimer could potentially function in some way as a signaling molecule (since the heterodimer would contain one wild-type extracellular domain and a wild-type cytoplasmic domain, albeit on different molecules).

We further examined the reduction of Diap1 protein in cells with reduced *ed* function. This finding is contrary to a previous study that reported that *ed* tissue had increased expression of Yki-target genes including *diap1* (Yue *et al*. 2012). In addition to finding that clones of cells with reduced *ed* function had reduced Diap1 protein levels (Figure 6F, F’), we also found that wild-type cells immediately adjacent to these clones sometimes had increased Diap1 protein levels - a “border effect” (Figure 4G, G’; Figure 6F, F’). We did not observe a change in *diap1* RNA either in the clone or in the adjacent cells by RNA in-situ hybridization using the hybridization chain reaction (data not shown). However, when we used a transcriptional reporter of *diap1* which contains 8 copies of a Hippo-responsive element (HRE) (Wu *et al*. 2008), we observed decreased expression in clones expressing *diap1-RNAi* and sometimes observed increased expression in cells just outside the clone (Figure 6G, G’). Also, contrary to that same study, we did not find elevated levels of nuclear Yki. Using a GFP-tagged Yki, which has proven to be a reliable indicator of Yki localization (Fletcher et al., 2018), we observed no change in the localization of Yki in *ed*-*RNAi* clones (Figure 6H, H’). Additionally, two other reporter of Yki-driven expression, *bantam-lacZ* (Herranz *et al*. 2012) (a direct reporter of bantam transcription, *not* the *bantam* sponge which is inversely correlated to *ban* levels) and *fj-lacZ* (Brodsky and Steller 1996), are expressed at lower levels in the clone and at higher levels in some neighboring cells (Figure 6I, I’, J, J’). In contrast to all of these reporters, another Yki-dependent reporter, *ex-lacZ* (Boedigheimer and Laughon 1993), was expressed at higher levels in cells expressing *ed-RNAi* (Figure 6K, K’). In summary, our results are inconsistent with the previous conclusion that *ed* mutant tissue has increased levels of nuclear Yki and increased expression of Yki-target genes. We observed no change in Yki localization and lower levels of expression of three Yki-dependent reporter genes. One, *ex-lacZ,* was indeed expressed at higher levels in *ed* mutant cells. When *ed* was overexpressed using *nub-Gal4*, there was a strong reduction in wing size (Supplementary Figure 2A, B). Once again, inconsistent effects on Yki-targets were observed. Diap1 protein levels were unaffected in *ed*-overexpressing clones (Supplementary Figure 2C, C’) but reduced when *ed* was overexpressed in the entire posterior compartment (Supplementary Figure 2D, D’). *ban-lacZ* expression and *ex-lacZ* expression were both increased (Supplementary Figure 2E, E’, F, F’). Taken together, these results do not support a simple relationship between Ed levels and Yki-target gene expression.

Border effects such as that observed in *ed* clones have been described previously with manipulations of the Fat/Dachsous pathway (Willecke *et al*. 2008; Matakatsu and Blair 2012). To investigate a possible role for this pathway, we stained discs containing *ed-RNAi* clones with anti-Fat. We observed increased staining in the clones (Figure 7A) which might be due to a combination of increased apical Fat levels as well as the reduced apical profiles of cells in the clone which has been noticed previously (Wei *et al*. 2005; Laplante and Nilson 2006; Chang *et al*. 2011). Importantly, we observed decreased anti-Fat staining in the cells immediately adjacent to the clone. A reduction in Fat levels would be expected to result in reduced Hippo pathway activity (Cho *et al*. 2006; Bennett and Harvey 2006; Silva *et al*. 2006; Willecke *et al*. 2006) and increased expression of Yki-target genes as we have observed.

**Figure 7:**
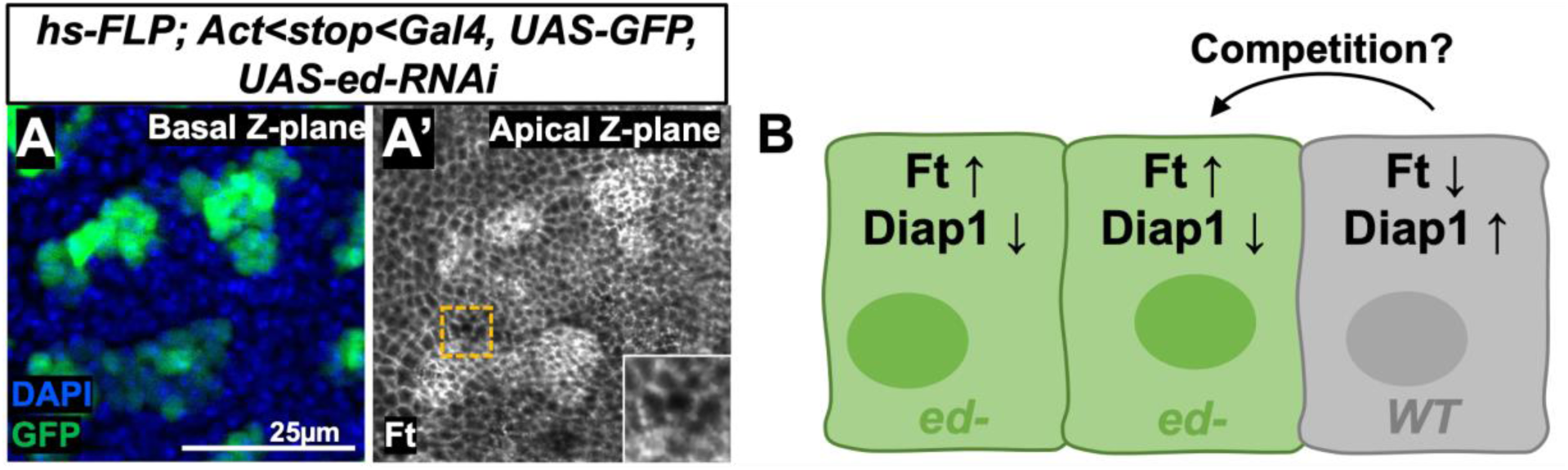
*echinoid* mutant cells could be eliminated by a mechanism similar to cell competition at clonal boundaries, (**A, A’**) GFP-marked clones expressing *ed-RNAi* (**A**) stained with an anti-Fat antibody. The inset shows a close-up of the boundary of the clone. (**B**) A model figure showing the effect of *ed* loss in *ed* cells and wild-type neighbors. Within *ed* clones, Fat levels are increased and Diap1 levels are decreased; the opposite is true in the wild-type neighbors. These differences may confer a competitive advantage to the wild-type neighbors, which could facilitate the elimination of *ed* clones from mosaic tissue.

## Discussion

In order to systematically assess the requirement of individual cell-surface proteins in regulating cell survival or proliferation in clones, we examined the effect of reducing the expression of 75 such genes that have detectable levels of expression in the wing disc. Although we did not verify the efficacy of the knockdown by RNAi in each case, we found that for 66 of these, there was no obvious decrease in clone size or alteration in clone shape. Thus, most cell surface proteins, at least individually, do not function to sustain cell survival or proliferation. We found that both reducing levels of either *ft* or *ds* resulted in rounder clones that appeared larger. Since these genes have been studied in extensive detail, we did not pursue them further.

Of the remaining genes identified in the screen, we chose to focus on *ed* for two reasons. First, reducing *ed* levels has very different effects in clones and in entire tissues. Clones with reduced *ed* function are eliminated, while imaginal discs that lack *ed* function in large regions are overgrown. In some ways this phenotype is reminiscent of cells that have disruptions in apicobasal polarity such as *scrib* (Bilder *et al*. 2000; Brumby and Richardson 2003; Hariharan and Bilder 2006) with the important difference being that cells lacking *ed* still seem to have preserved most aspects of their apicobasal polarity, and tissue architecture appears relatively normal.

Second, previous studies have reported that *ed* mutant tissue has increased EGFR signaling (Bai *et al*. 2001; Islam *et al*. 2003; Rawlins *et al*. 2003a; Spencer and Cagan 2003; Fetting *et al*. 2009; Ho *et al*. 2010) and reduced activity of the Hippo pathway resulting in increased Yki-target gene expression (Yue *et al*. 2012). Both of these changes would be expected to promote cell survival and proliferation. Based on these observations, the underrepresentation and elimination of *ed* mutant clones in imaginal disc is unexpected. Previous work, however, has also shown that *ed* mutant cells appear to sort away from wild-type cells and that there is an actomyosin cable generated at the clone interface in wild-type cells that could potentially promote clone extrusion. We confirmed that EGFR signaling is likely elevated as assessed by reduced levels of nuclear Capicua (Supplementary Figure 3A-C) and also observed an actomyosin cable at the clone borders (Supplementary Figure 3D) like others have reported (Wei *et al*. 2005; Laplante and Nilson 2006, 2011; Lin *et al*. 2007; Chang *et al*. 2011). However, our observations are not consistent with the proposed effect on the Hippo pathway (Figure 6E-J, Supplementary Figure 2C-F).

### Why are *ed* clones eliminated?

We have found that *ed* mutant tissue has reduced levels of the anti-apoptotic protein Diap1. We observed this consistently in *ed* clones as well as in the entire posterior compartment of the wing disc when *ed* function was reduced using RNAi. Furthermore, the elimination of *ed* clones could be rescued by co-expressing *diap1* with *ed-RNAi*. Together, these observations suggest the reduction in Diap1 levels underlies, in significant part, the elimination of *ed* clones.

Although we could not detect a reduction in *diap1* RNA by *in situ* hybridization (data not shown), we were able to detect decreased expression of a commonly-used *diap1* transcriptional reporter (Wu *et al*. 2008) which is a more sensitive reagent. Since this reporter was constructed using multiple copies of a Hippo-responsive element from the *diap1* locus, these data indicate that the change in Diap1 protein levels is due, at least in part, to a decrease in Yki-dependent *diap1* transcription. This finding differs from a previous study that reported increased Diap1 expression in *ed* clones that was attributed to increased expression of Yki target genes (Yue *et al*. 2012). In contrast to the conclusions of that study, we find decreased expression of at least three different transcriptional reporters of Yki in *ed* mutant clones: *diap-lacZ*, *ban-lacZ* and *fj-lacZ* and also no change in nuclear Yki localization. We do, however, find that a different reporter of Yki-dependent transcription, *ex*-*lacZ* is expressed at increased levels in *ed* mutant tissue indicating that the regulation of Yki-target genes by Ed is not easily explained by a simple effect on alterations in Hippo pathway activity.

Ed depleted tissues have increased apoptosis. We observed this in clones, especially at clone boundaries, and when *ed* was depleted in broader domains. Our results also explain why a reduction of *ed* function in the posterior compartment of the wing disc slows its growth; the affected posterior compartments are smaller than equivalently staged wild-type discs. This could simply be because the increased level of cell death in this compartment reduces tissue mass.

We have also identified a boundary effect at the border of *ed* clones: Some, but not all, wild-type cells immediately adjacent to the clones have elevated expression of Yki-target genes and elevated Diap1 protein. In this way, these cells resemble cells that have mutations in the Hippo pathway and behave as supercompetitors (Tyler *et al*. 2007). Thus, the reduction of Diap1 in *ed* mutant clones coupled with the generation of boundary cells that have some of the properties of supercompetitors could together contribute to the elimination of *ed* cells from the disc (Figure 7B).

### Why are *ed* mutant tissues overgrown?

Although *ed* tissue grows more slowly than wild-type tissue, it appears to grow beyond an appropriate size. Thus, Ed has a second function in sensing tissue size and arresting growth at the appropriate time. Although not definitive, our experiments suggest that this function of Ed might primarily involve its role as an adhesion molecule. The levels of E-cadherin are known to be altered in *ed* mutant tissue and Ed and E-cadherin may function with some degree of redundancy to mediate cell-cell adhesion (Wei *et al*. 2005; Laplante and Nilson 2006; Chang *et al*. 2011). The mechanisms by which tissues sense their final size and arrest their growth accordingly are still poorly understood. While growth arrest is clearly disrupted in mutants that completely disrupt apicobasal polarity (Hariharan and Bilder 2006), it is possible that even subtle changes in junctional components could disrupt growth arrest.

### Concluding remarks

Ed participates in, and regulates, at least two key cellular processes: adhesion at the adherens junction (Wei *et al*. 2005; Laplante and Nilson 2006, 2011; Chang *et al*. 2011), and endocytosis and endosomal trafficking (Rawlins *et al*. 2003b; Fetting *et al*. 2009; Ho *et al*. 2010; Li *et al*. 2015; Yang *et al*. 2018; Chan *et al*. 2021). Moreover, at least based on genetic interactions, *ed* influences the EGFR, Notch, and Hippo pathways. Ed may interact with more pathways or processes than we currently appreciate, either directly or indirectly via its effects on adhesion, endomembrane trafficking, or through signaling crosstalk. The collection of phenotypic abnormalities and alterations in gene expression that we observe in *ed* mutants are therefore likely to involve a summation of alterations in multiple pathways rather than a disruptive effect on any single pathway.

## Methods

### Fly Stocks & Husbandry

Unless otherwise noted, all experimental crosses were raised at 25°C on food prepared according to the recipe from the Bloomington *Drosophila* Stock Center.

Stocks used in this study include or were derived from the following: *Oregon-R* (“Ore-R,” used as wild type), *y w hs-FLP; act<[y+]<Gal4 UAS-GFP/SM5-TM6B*, *hs-FLP;; act<stop<Gal4 UAS-RFP/SM5-TM6B*, *TIE-DYE* (Worley *et al*. 2013), *FRT40A* and *FRT40A, white+ ubi-GFP* (Xu and Rubin 1993), *FRT40A MARCM* (Lee and Luo 1999), *eyFLP; FRT40A CL white+/CyO* (BL5622), *UAS-ed* (Bai *et al*. 2001), *ed^IF20^ FRT40A*, *ed^1×5^ FRT40A*, and *ed^sIH8^ FRT40A* (Bai *et al*. 2001), *UAS-ed^Full^*and *UAS-ed^ΔC^-GFP* (Laplante and Nilson 2011), *Yki:GFP* (Fletcher et al., 2018), *nub-Gal4* (*AC-62,* BL25754), *hh-Gal4* (BL45169), *ban-lacZ* (BL10154, Herranz et al., 2012), *fj-LacZ* (*fj^P1^*, BL44253; Brodsky & Steller, 1996), *ex-lacZ* (Boedigheimer and Laughon 1993), *diap1-GFP.HREx8* (Wu *et al*. 2008)*, UAS-w-RNAi* (BL33644), *UAS-ama-RNAi* (BL33416), *UAS-beat-Vc-RNAi* (BL60067), *UAS-Cont-RNAi* (BL34867), *UAS-shg-RNAi* (BL32904), *UAS-side-VII-RNAi* (V10011), *UAS-ft-RNAi* (BL34970), *UAS-ds-RNAi* (BL32964), *UAS-ed-RNAi* (V104279, V3087, V938, BL38423 [“*ed-RNAi*” refers to V104279 unless otherwise indicated]), *UAS-otk2-RNAi* (BL55892), *UAS-p35* (BL5073). “BL#” or “V#” indicates stocks obtained from the Bloomington *Drosophila* Stock Center (BDSC; Bloomington, IN, USA) or Vienna *Drosophila* Resource Center (VDRC; Vienna, Austria), respectively.

Additional stocks which were included in the genetic screen but are not mentioned in the main text are listed in Supplementary Table 3 with BDSC or VDRC numbers indicated.

### Mosaic tissue generation

Clones induced by heat shock were generated in a 37°C water bath 48 h before dissection, unless otherwise noted. FLP-out Gal4 clones were made using heat shocks of 12 minutes (to generate clones at low density), 15 minutes (for medium density), or 30 minutes (for high density). MARCM clones and mitotic recombination clones were generated using a 1-hr heat shock.

Mitotic recombination clones made in the eye were induced by expression of the *eyFLP* driver.

### Screen

∼10 *UAS-RNAi* males were crossed to ∼20 *y, w, hs-FLP; act<[y+]<Gal4, UAS-GFP/SM5-TM6B or TIE-DYE (Act<stop<lacZ.nls, Ubi <stop<GFP.nls; Act<stop<GAL4, UAS-his2A::RFP/SM5-TM6B)* virgin females. Crosses were kept on Bloomington food supplemented with yeast and flipped once daily. Clones were induced by heat shock on day 3. Early rounds of screening used a 15-minute heat shock, although we later switched to a 12-minute heat shock for the majority of the screen since the low clone density made identifying deviations in either direction easier. Wing imaginal discs from ∼6 wandering L3 larvae per line were dissected ∼48h after heat shock, stained with DAPI, and imaged.

### Immunohistochemistry and fluorescence microscopy

Imaginal discs were dissected in PBS, fixed in 4% paraformaldehyde in PBS, and permeabilized in 0.1% Triton in PBS. Primary antibody incubations were done overnight at 4°C. Secondary antibody incubations were done for 2-3h at room temperature, or overnight at 4°C. Discs were mounted in SlowFade Diamond Antifade Mountant (S36963, Invitrogen).

Primary antibodies used are: rabbit anti-Ed (1:500, J-C. Hsu, Wei et al., 2005), rabbit anti-cleaved Dcp-1 (1:250; Asp216, Cell Signaling Technology), mouse anti-Diap1 (1:200, B. Hay), mouse anti-β-Galactosidase (1:500, WH0051083M1 Sigma-Aldrich), mouse anti-β-Galactosidase (1:500, SAB4200805, Sigma-Aldrich), chicken anti-GFP (1:500, ab13970 Abcam); rat anti-Ci (1:500, #2A1; Developmental Studies Hybridoma Bank, DSHB), rat anti-Fat (1:400, K. Irvine, Feng & Irvine, 2009), and guinea pig anti-Cic (1:300, Tseng et al., 2007).

Secondary Alexa-Fluor antibodies from Invitrogen were used at 1:500. Nuclei were stained with DAPI (1:1,000, Cell Signaling).

Fluorescence images were taken on a Zeiss Axio Imager M2 equipped with a 20× objective (Plan-Apochromat, 20×/0.8, Zeiss), LED light source (Excelitas Technologies), AxioCam 506 mono camera (Zeiss), and ApoTome.2 slider for optical sectioning. Images and image stacks were acquired and optically sectioned in ZEN 2.3 software (Zeiss). Imaged were processed using FIJI software (Schindelin *et al*. 2012). Unless otherwise noted, images show a single Z-plane.

### Adult wing imaging and quantification

Adult wings were dissected from female flies. One wing per fly was mounted in Gary’s Magic Mountant (Lawrence *et al*. 1986). Wings were imaged using a Keyence VHX-5000 digital microscope, using the 20-200× lens at 150×. Brightness, contrast, and color tone of wing images have been adjusted on some images for improved visibility of features relevant to this study (wing shape and size).

For qualitative comparisons of wing sizes, wing images or traced silhouettes were overlaid in PowerPoint. For quantitative comparisons of wing sizes, wing outlines were traced and area was quantified in Fiji. Charts were generated using the ggplot2 package in RStudio (Wickham 2016).

Wing aspect ratios were calculated by dividing the length of proximodistal (PD) axis (measured from the posterior junction of the wing and hinge to the tip of the L3 vein) by the length of the anteroposterior (AP) axis (measured as the shortest distance from the tip of the L5 vein to the L1 margin).

### Statistical analysis

P values were obtained by one-way ANOVA with Tukey’s HSD test using Astatsa freeware (Vasavada 2016). P value significance <0.01: **; 0.01 to 0.05: *; >0.05: not significant. All error bars show standard deviation.

### Hybridization Chain Reaction

*In situ* hybridization chain reaction (HCR) was performed on wing discs based on HCR v3.0 protocol (Choi *et al*. 2018; S Bruce *et al*. 2021) excluding methanol dehydration. Larvae were dissected, fixed in 4% paraformaldehyde in PBS, permeabilized, and incubated overnight with RNA probes at 37°C. Samples were then washed and incubated overnight with fluorescently-tagged RNA hairpins and DAPI at room temperature. Probe sequences were designed in an open-source probe design program (ÖzpolatLab-HCR 2021) and synthesized by Integrated DNA Technologies. Hairpins and buffers were from Molecular Instruments.

## Acknowledgements

We thank current and former members of the Hariharan laboratory for helpful feedback and discussions, especially Luigi Viggiano, who conducted related experiments that influenced our thinking on this study, and Melanie Worley. We also thank Bruce Hay, Jui-Chou Hsu, Ken Irvine, Laura Nilson, Duojia Pan, Nic Tapon, the Bloomington Drosophila Stock Center, the Vienna Drosophila Resource Center, and the Developmental Studies Hybridoma Bank for reagents and fly stocks. This work was funded by a NIH grant R35 GM122490 (to I.K.H) and NSF GRFP DGE 1752814 (to D.C.S).

## Author Contributions

DCS and IKH conceptualized and designed the study. DCS, WYS, and AR-V performed the experiments. DCS conducted data analysis and visualization. DCS and IKH wrote the manuscript. All authors read and approved the manuscript.

---

**Supplementary Figure 1:**
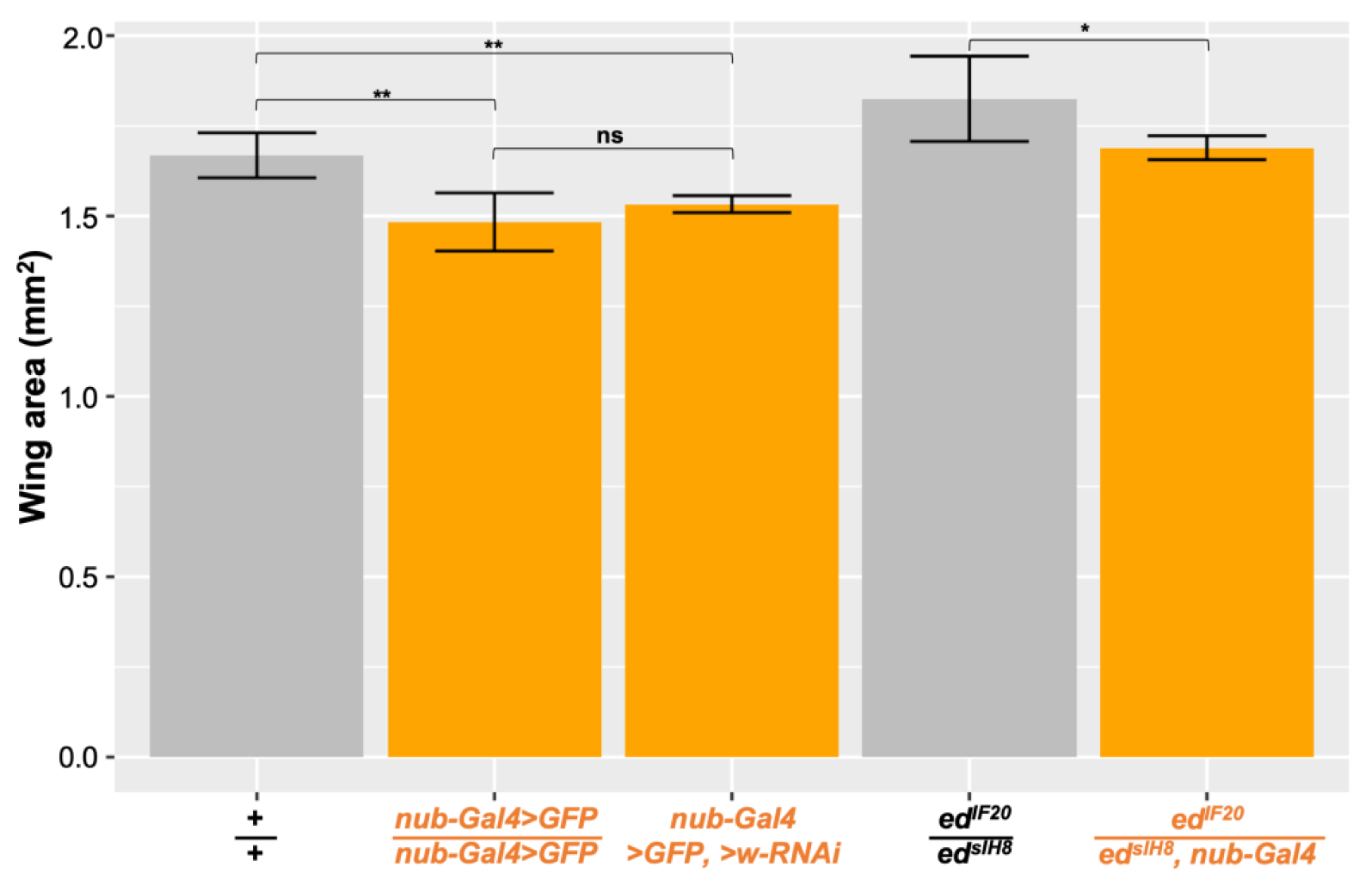
Effect of the *nub-Gal4* chromosome on wing size. Effect of *nub-Gal4* on size of adult wings. Quantification of wing areas of “wild type” (*Oregon-R* or *nub-Gal4* driving UAS constructs [*GFP, w-RNAi*] that should not affect sing size) and *ed*-depleted wings, with and without *nub-Gal4.* Wing blades are smaller in flies carrying the *nub-Gal4* chromosome. n=10 wings (*+/+* [*Oregon-R*]*; ed^sIH8^/ed^IF20^; nub-Gal4, >GFP, >w-RNAi;* same as shown in Figure 5I, 6D), 11 wings (*nub-Gal4, >GFP* [homozygotes]), 4 wings (*ed^sIH8^/ed^IF20^, nub-Gal4;* same as Figure 6D). “ns” indicates p> 0.05, ** indicates p< 0.01, calculated using ANOVA with post-hoc Tukey’s HSD test. Error bars indicate standard deviation.

**Supplementary Figure 2:**
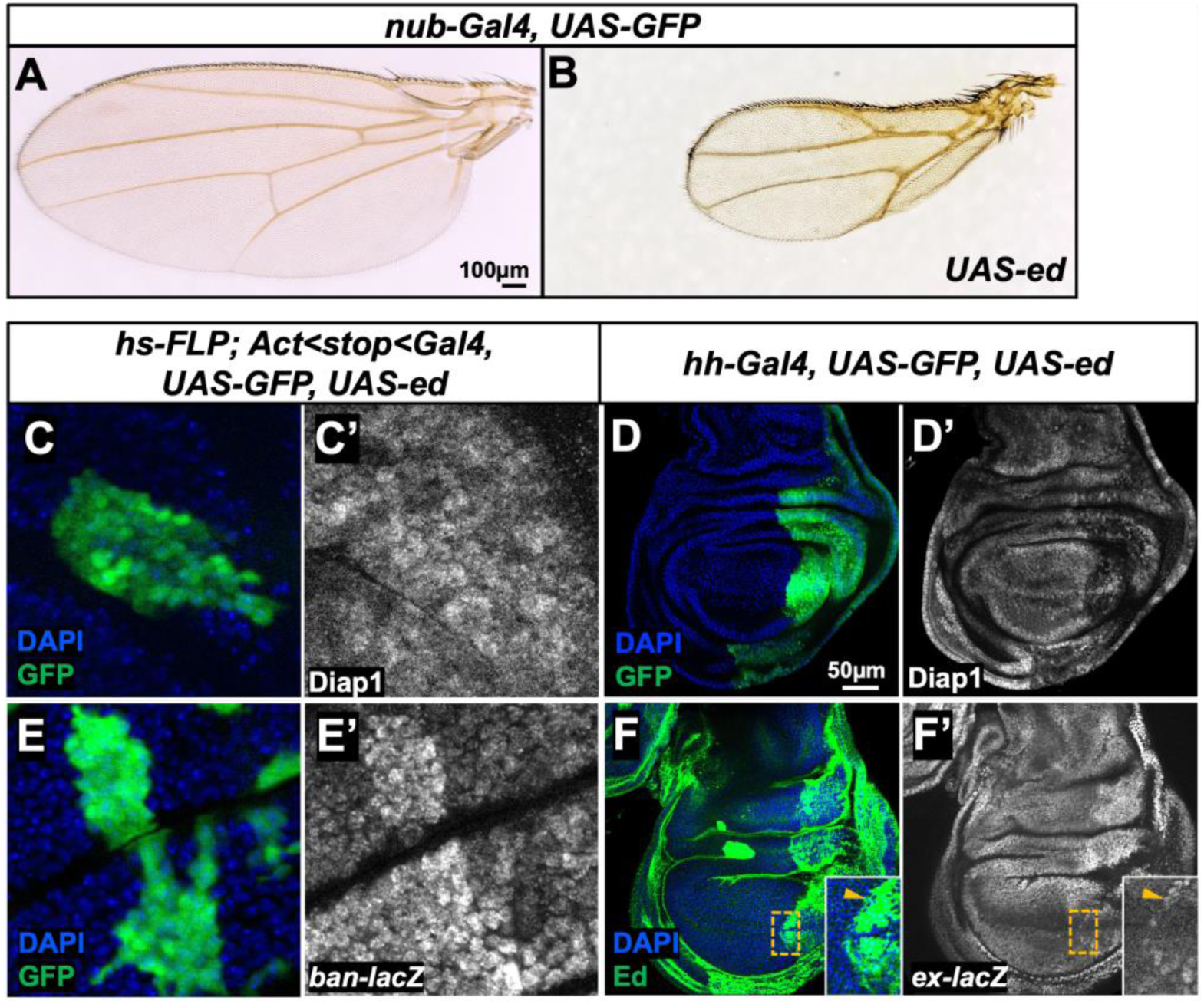
Phenotypes resulting from *echinoid* overexpression. (**A, B**) Effect of *ed* overexpression on wing size. Panels show adult wings of the indicated genotype. Wings overexpressing *ed* (**B**) are severely reduced in size, especially in the proximal regions. (**C, D**) Effect of *ed* overexpression on Diap1 levels. GFP-marked clone that expresses *UAS-ed* (**C**) showing no obvious effect on Diap1 levels (**C’**). Overexpression of *ed* in the entire posterior compartment (**D**) results in a decrease in Diap1 (**D’**). (**E**) Overexpression of *ed* in GFP-marked clones (**E**) results in an increase in *ban-lacZ* expression (**E’**). (**F**) Overexpression of *ed* in the posterior compartment results in increased *ex-lacZ* expression (**F’**). The autonomous increase is less obvious in the pouch, except in the *ed*-overexpressing cells at the border with wild-type cells (inset, arrowhead).

**Supplementary Figure 3:**
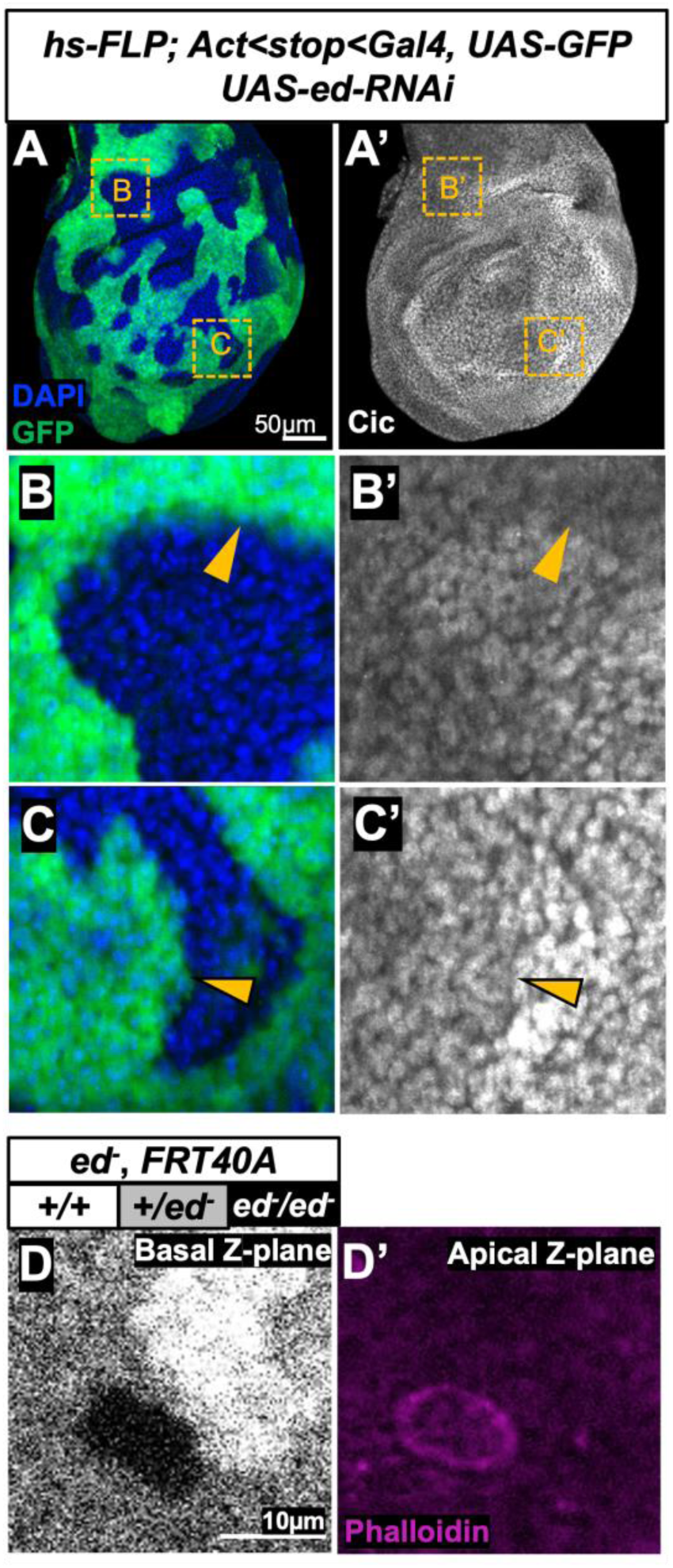
Other phenotypes observed with *echinoid* clones. (**A-C**) Effect of *ed* knockdown on Capicua (Cic) levels. (**A**) Cic levels are lower in some *ed* clones (arrowheads) than in wild-type neighbors, indicating increased EGFR activity within the clones. (**B, C**) and (**B’, C’**) show GFP-marked clones and Cic antibody staining at higher magnification. The location of these clones is indicated in (**A, A’**) by dashed lines. (**D, D’**) An apical actomyosin cable is observed at the interface of *ed* mutant tissue and the wild type neighbors (*ed^IF20^/ ed^IF20^* mitotic recombination clone is shown), consistent with previous reports (Wei *et al.* 2005; Laplante and Nilson 2006, 2011; Lin *et al.* 2007; Chang *et al.* 2011).

**Supplementary Table 1:**
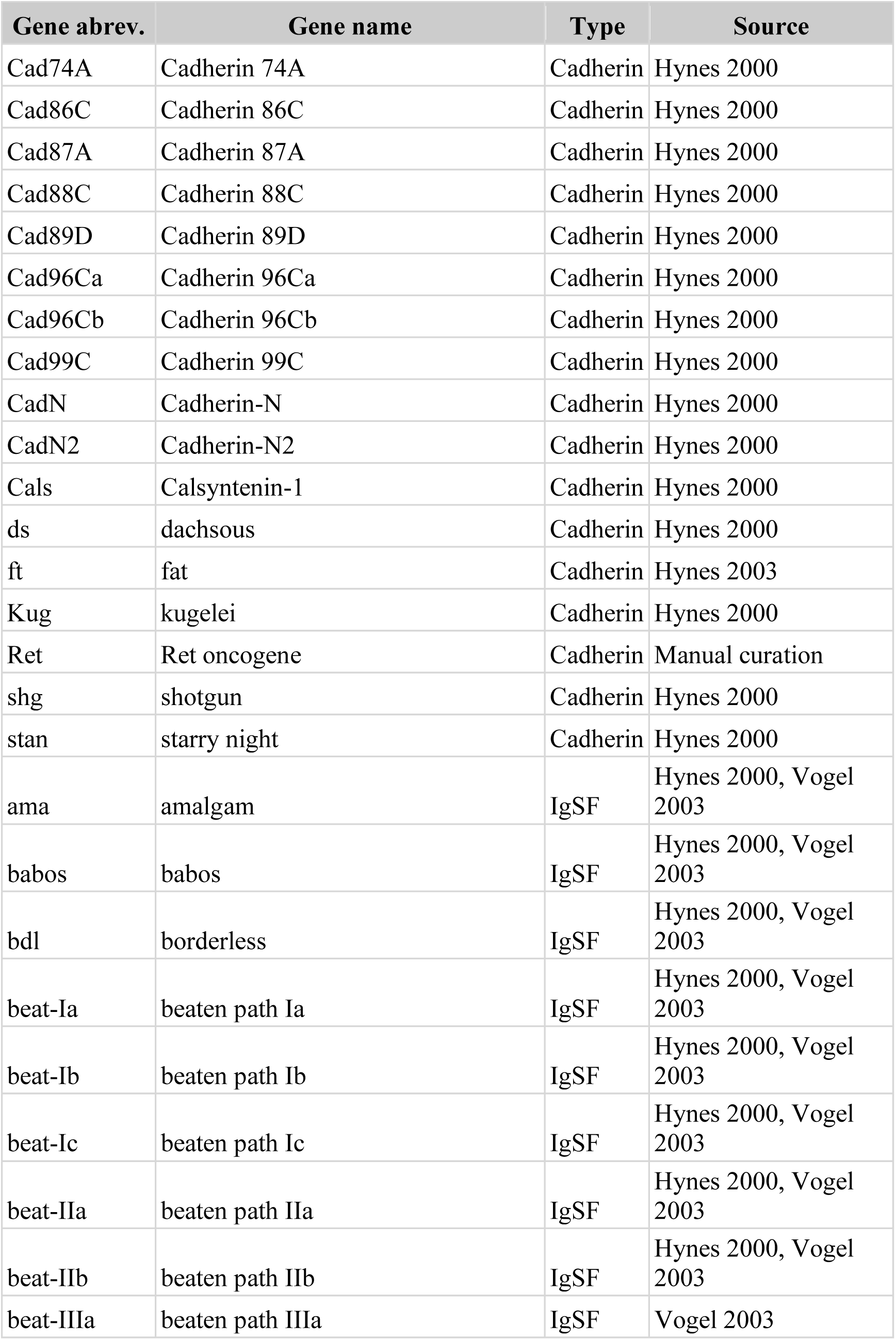
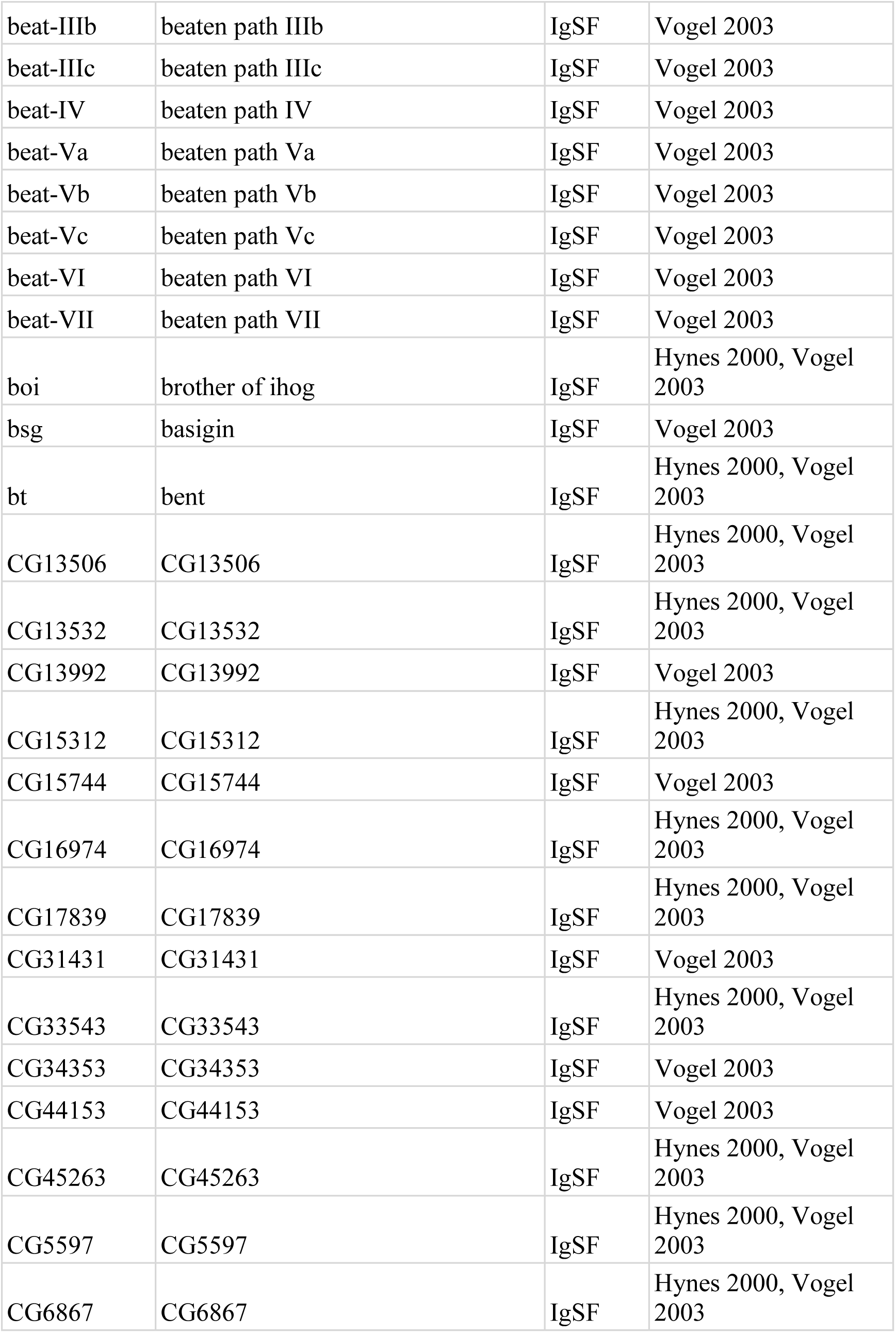
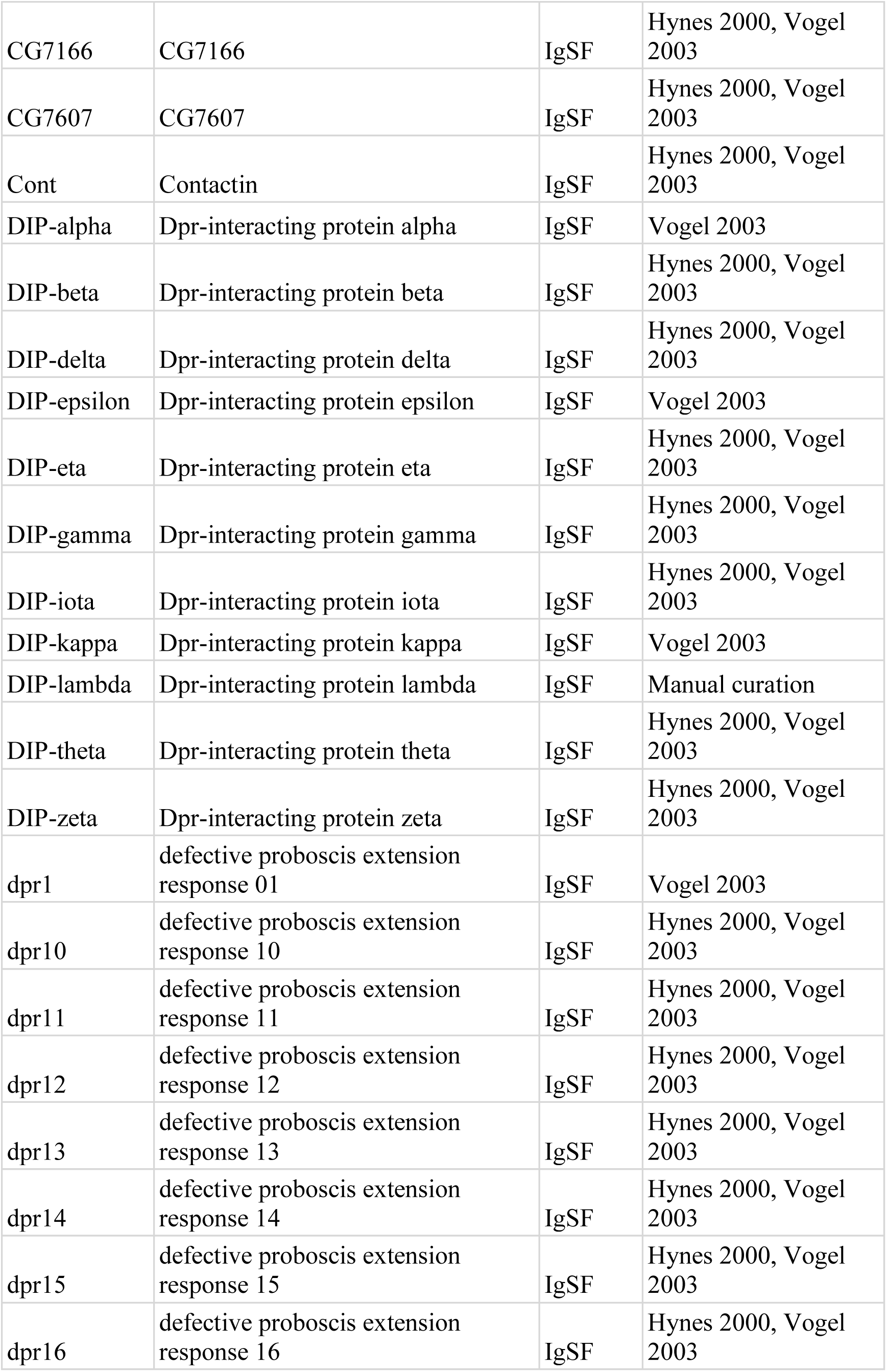
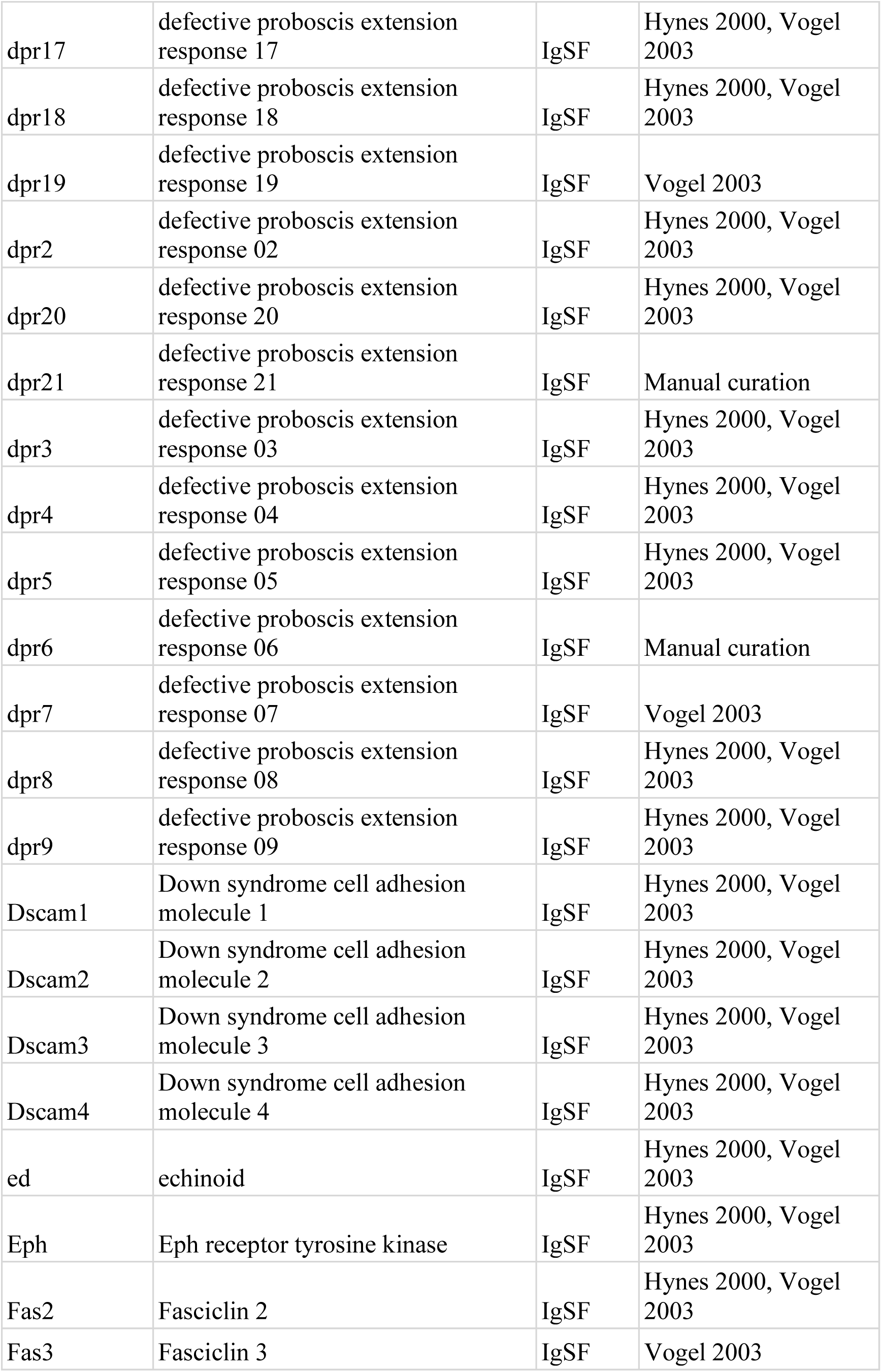
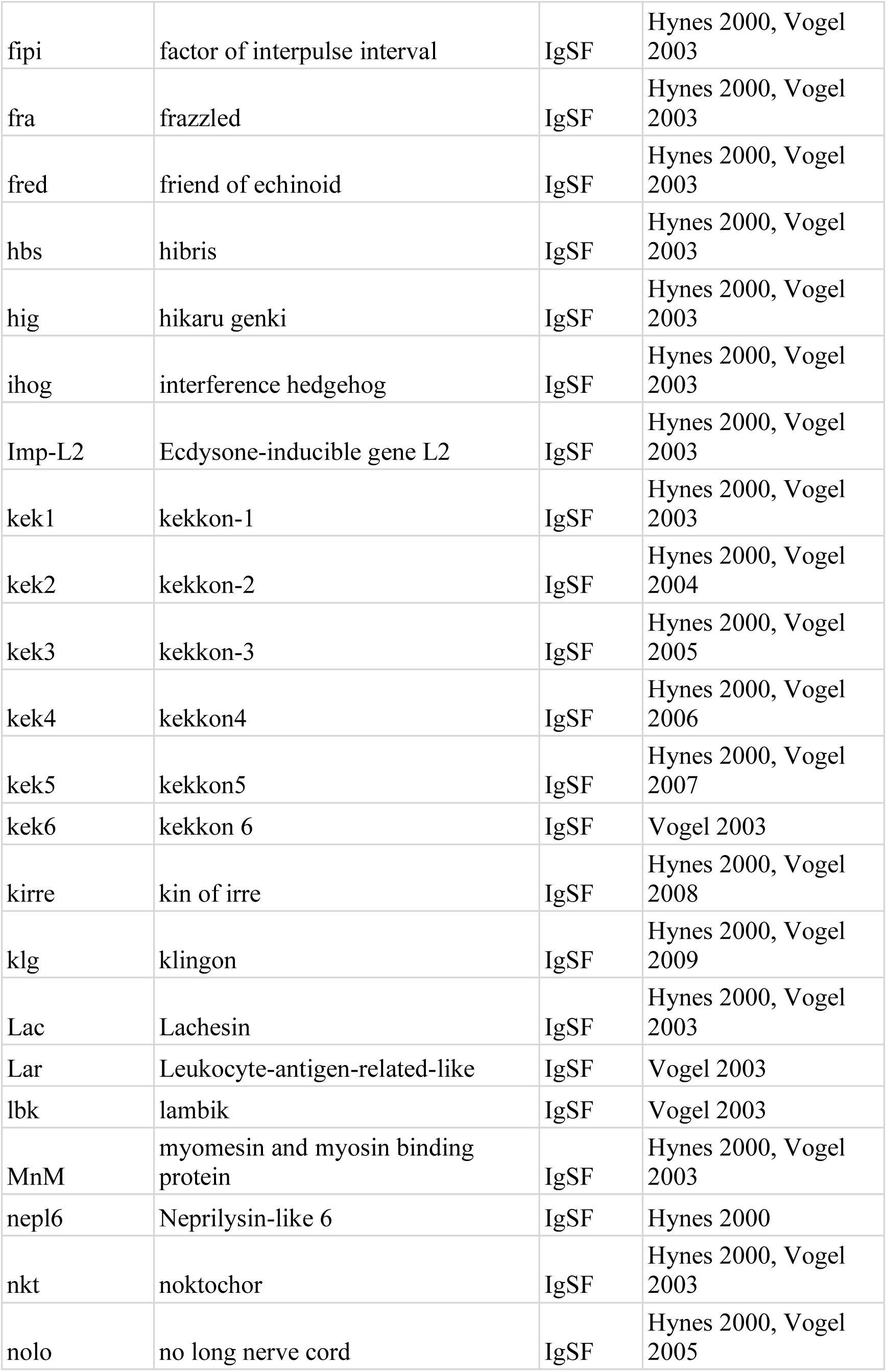
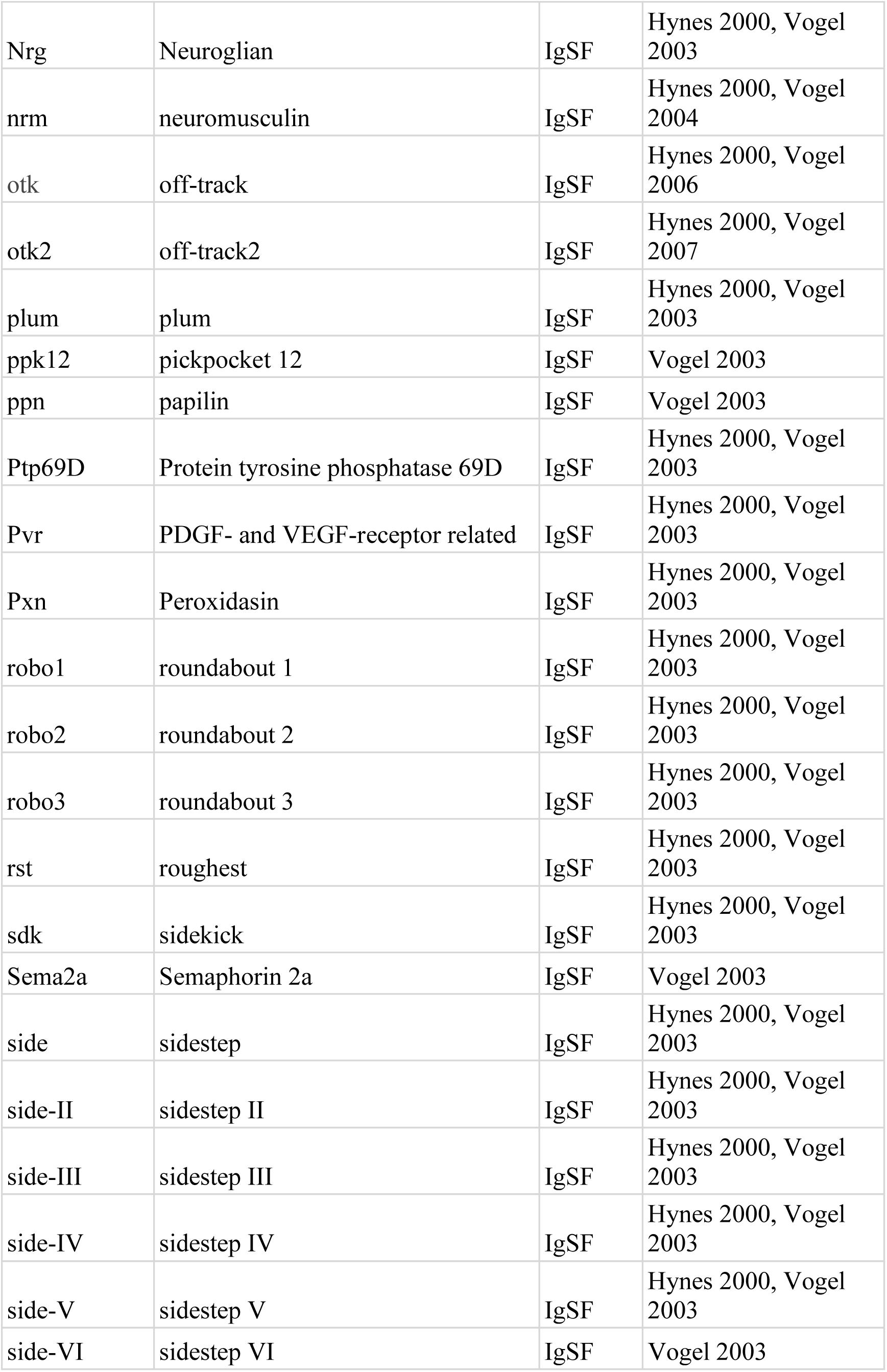
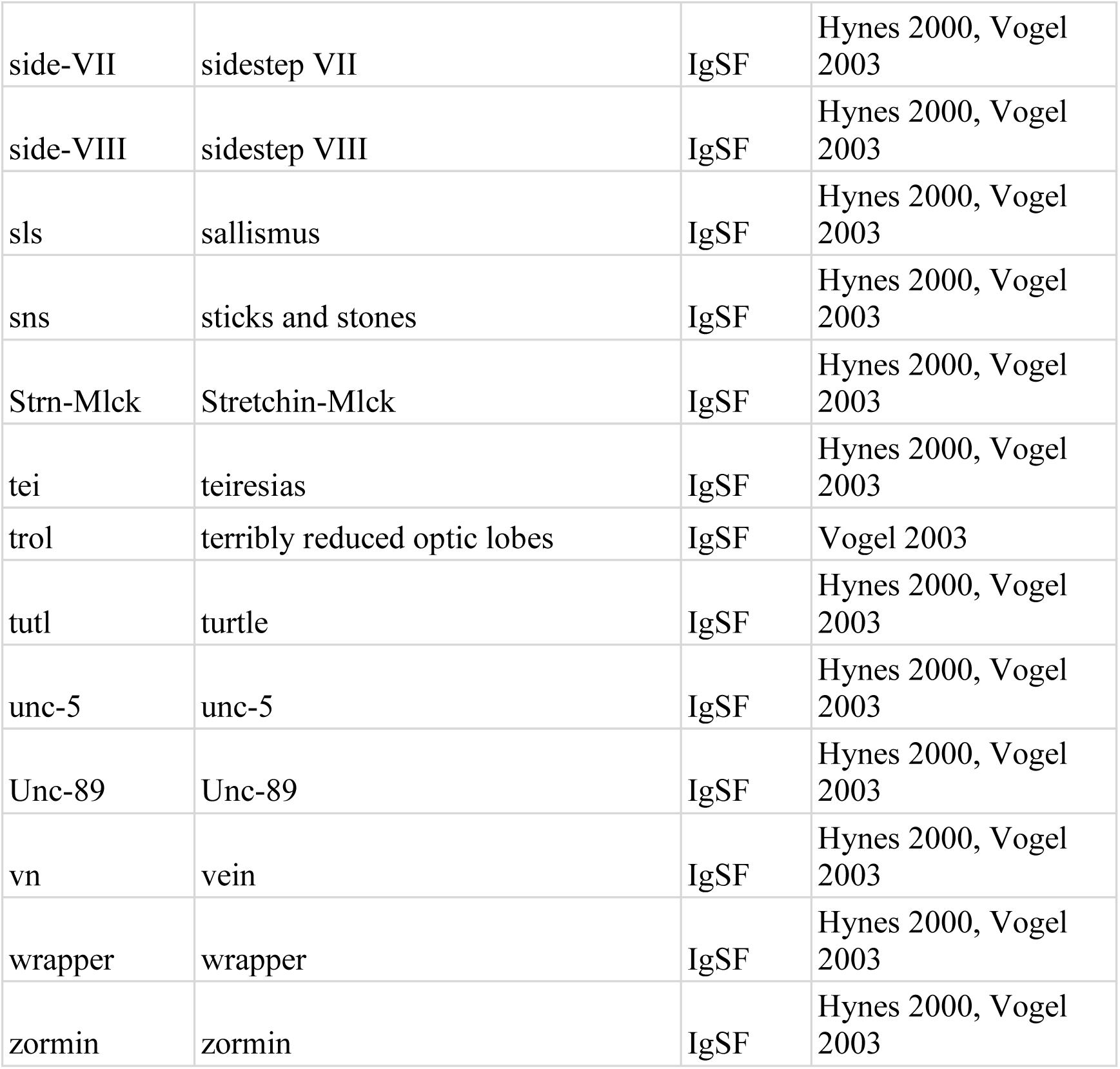
Initial candidate list.

**Supplementary Table 2:** Genes excluded from screen.

| **Gene abbrev.** | **Gene name** | **Criteria for exclusion** |
| --- | --- | --- |
| ppk12 | pickpocket 12 | Known to not be adhesion molecule (ENaC subunit); Incorrectly classified as IgSF? |
| htl | heartless | Known to not be adhesion molecule (RTK) |
| Ret | Ret oncogene | Known to not be adhesion molecule (RTK) |
| bt | bent | Known to not be adhesion molecule (sarcomere component) |
| sls | sallismus | Known to not be adhesion molecule (sarcomere component) |
| Unc-89 | Unc-89 | Known to not be adhesion molecule (sarcomere component) |
| zormin | zormin | Known to not be adhesion molecule (sarcomere component) |
| Sema2a | Semaphorin 2a | Known to not be adhesion molecule (secreted) |
| btl | breathless | Known to not be adhesion molecule (RTK) |
| beat-Ia | beaten path Ia | No expression RNAseq detected in wing disc |
| beat-Ib | beaten path Ib | No expression RNAseq detected in wing disc |
| beat-Ic | beaten path Ic | No expression RNAseq detected in wing disc |
| Cad88C | Cadherin 88C | No expression RNAseq detected in wing disc |
| Cad89D | Cadherin 89D | No expression RNAseq detected in wing disc |
| CadN | Cadherin-N | No expression RNAseq detected in wing disc |
| CadN2 | Cadherin-N2 | No expression RNAseq detected in wing disc |
| CG13532 | CG13532 | No expression RNAseq detected in wing disc |
| CG17839 | CG17839 | No expression RNAseq detected in wing disc |
| CG31431 | CG31431 | No expression RNAseq detected in wing disc |
| CG6867 | CG6867 | No expression RNAseq detected in wing disc |
| DIP-beta | Dpr-interacting protein beta | No expression RNAseq detected in wing disc |
| DIP-delta | Dpr-interacting protein delta | No expression RNAseq detected in wing disc |
| DIP-eta | Dpr-interacting protein eta | No expression RNAseq detected in wing disc |
| DIP-gamma | Dpr-interacting protein gamma | No expression RNAseq detected in wing disc |
| DIP-iota | Dpr-interacting protein<br>iota | No expression RNAseq detected in wing disc |
| DIP-theta | Dpr-interacting protein<br>theta | No expression RNAseq detected in wing disc |
| DIP-zeta | Dpr-interacting protein<br>zeta | No expression RNAseq detected in wing disc |
| dpr10 | defective proboscis<br>extension response 10 | No expression RNAseq detected in wing disc |
| dpr11 | defective proboscis<br>extension response 11 | No expression RNAseq detected in wing disc |
| dpr12 | defective proboscis<br>extension response 12 | No expression RNAseq detected in wing disc |
| dpr13 | defective proboscis<br>extension response 13 | No expression RNAseq detected in wing disc |
| dpr15 | defective proboscis<br>extension response 15 | No expression RNAseq detected in wing disc |
| dpr2 | defective proboscis<br>extension response 02 | No expression RNAseq detected in wing disc |
| dpr20 | defective proboscis<br>extension response 20 | No expression RNAseq detected in wing disc |
| dpr3 | defective proboscis<br>extension response 03 | No expression RNAseq detected in wing disc |
| dpr4 | defective proboscis<br>extension response 04 | No expression RNAseq detected in wing disc |
| dpr5 | defective proboscis<br>extension response 05 | No expression RNAseq detected in wing disc |
| dpr8 | defective proboscis<br>extension response 08 | No expression RNAseq detected in wing disc |
| Dscam2 | Down syndrome cell<br>adhesion molecule 2 | No expression RNAseq detected in wing disc |
| Dscam3 | Down syndrome cell<br>adhesion molecule 3 | No expression RNAseq detected in wing disc |
| Dscam4 | Down syndrome cell<br>adhesion molecule 4 | No expression RNAseq detected in wing disc |
| fipi | factor of interpulse<br>interval | No expression RNAseq detected in wing disc |
| kek3 | kekkon-3 | No expression RNAseq detected in wing disc |
| kek4 | kekkon4 | No expression RNAseq detected in wing disc |
| klg | klingon | No expression RNAseq detected in wing disc |
| nepl6 | Neprilysin-like 6 | No expression RNAseq detected in wing disc;<br>Incorrectly classified as IgSF? |
| nolo | no long nerve cord | No expression RNAseq detected in wing disc |
| robo3 | roundabout 3 | No expression RNAseq detected in wing disc |
| side-II | sidestep II | No expression RNAseq detected in wing disc |
| side-III | sidestep III | No expression RNAseq detected in wing disc |
| side-VI | sidestep VI | No expression RNAseq detected in wing disc |
| side-VIII | sidestep VIII | No expression RNAseq detected in wing disc |

**Supplementary Table 3:**
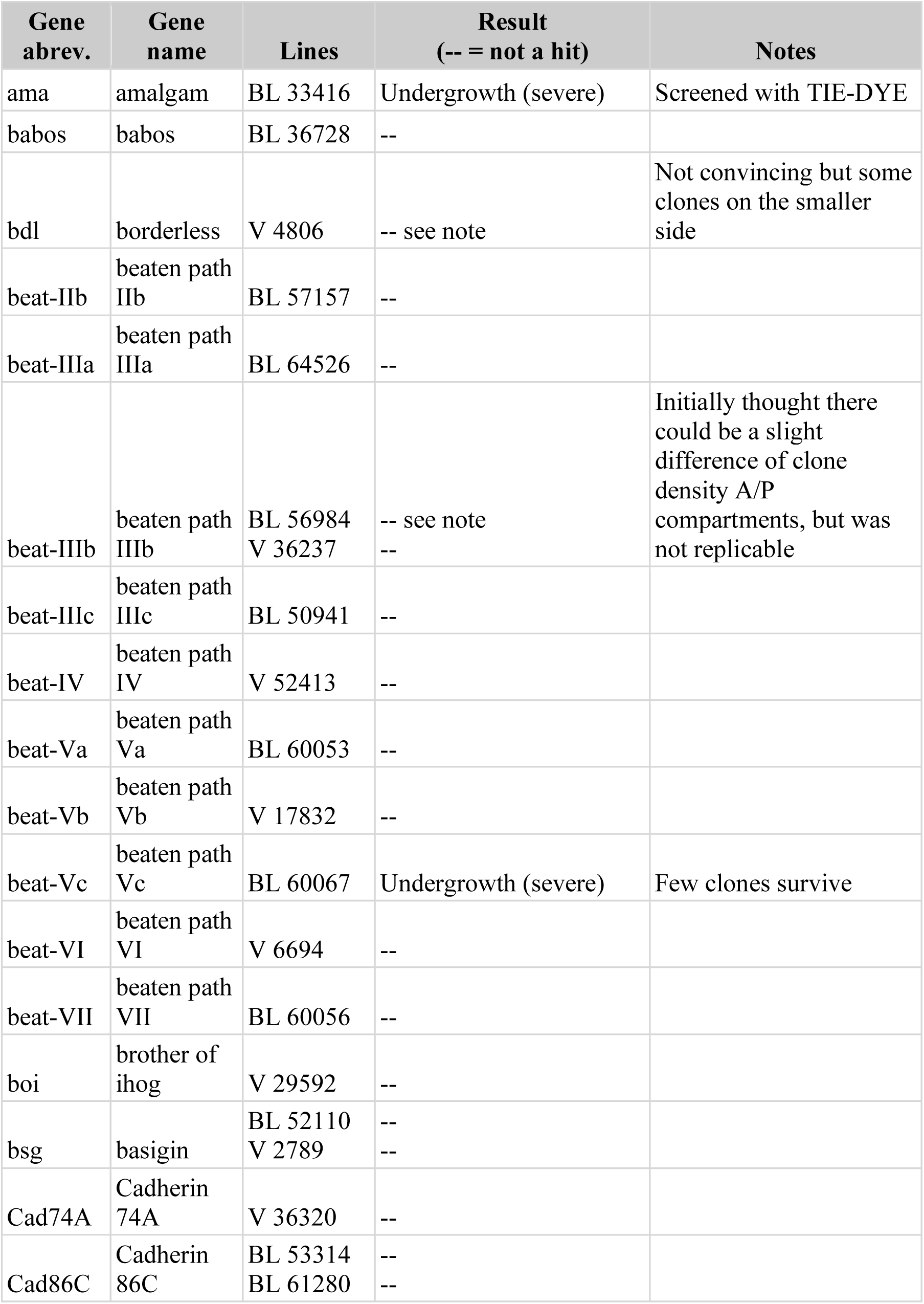
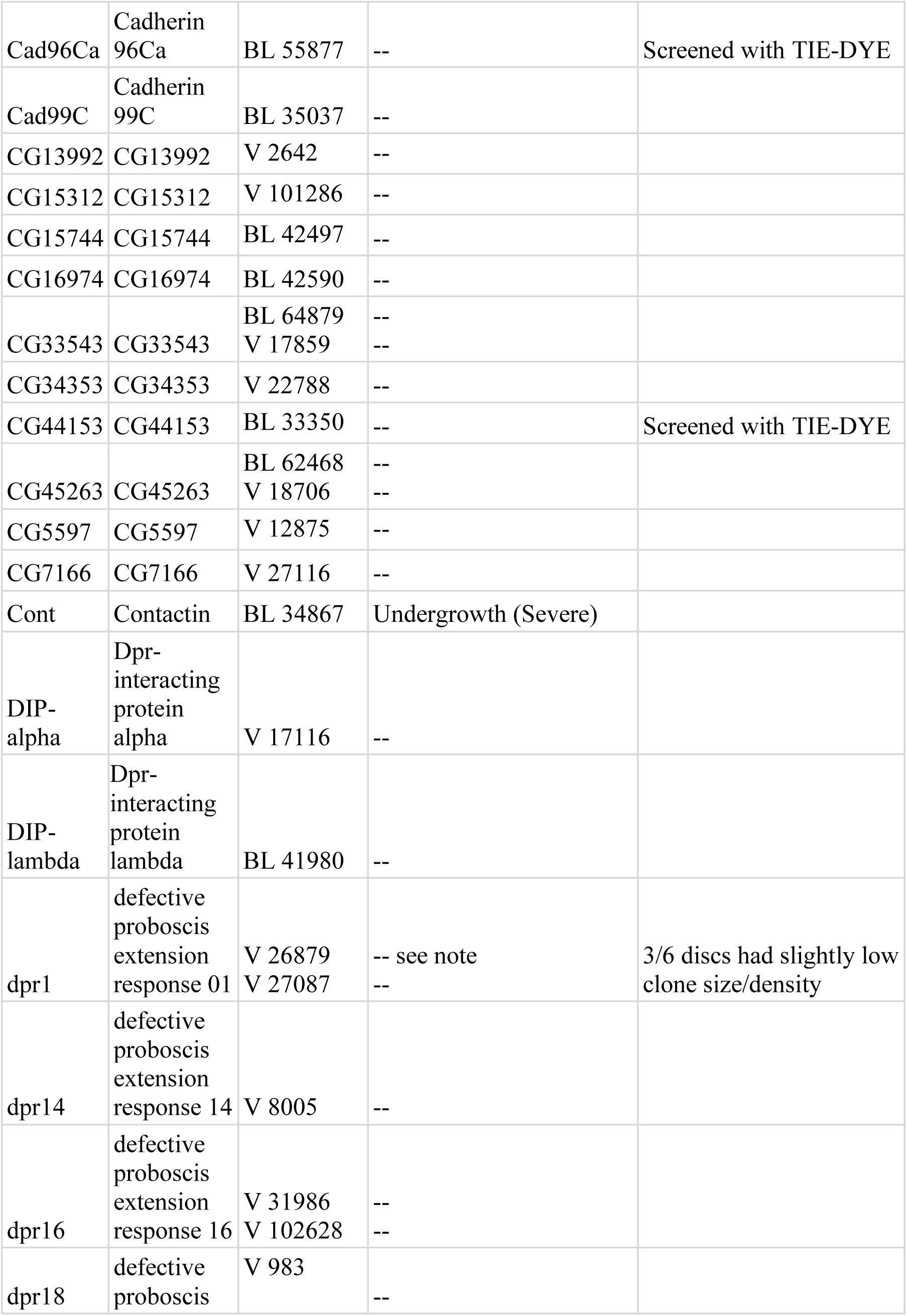
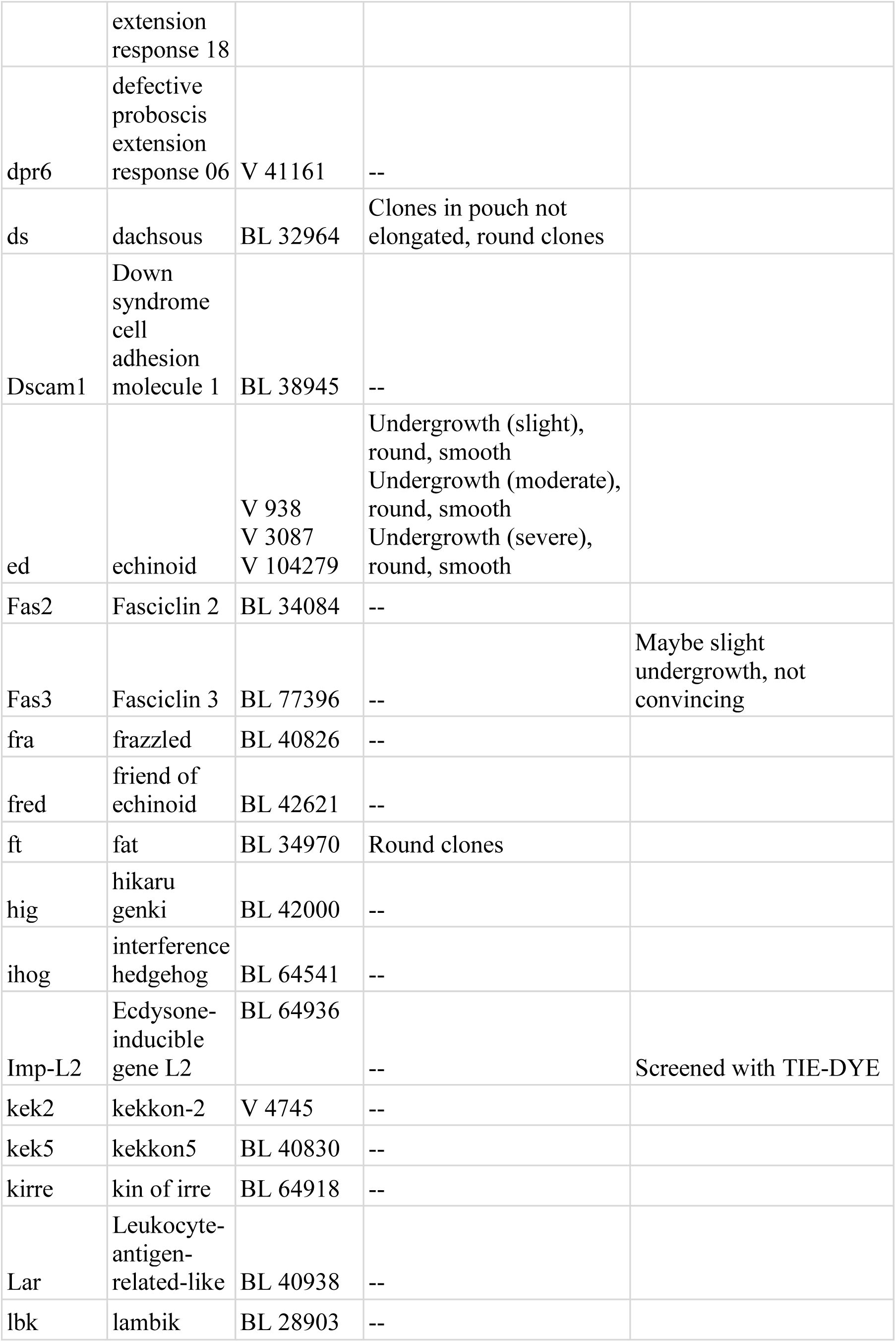
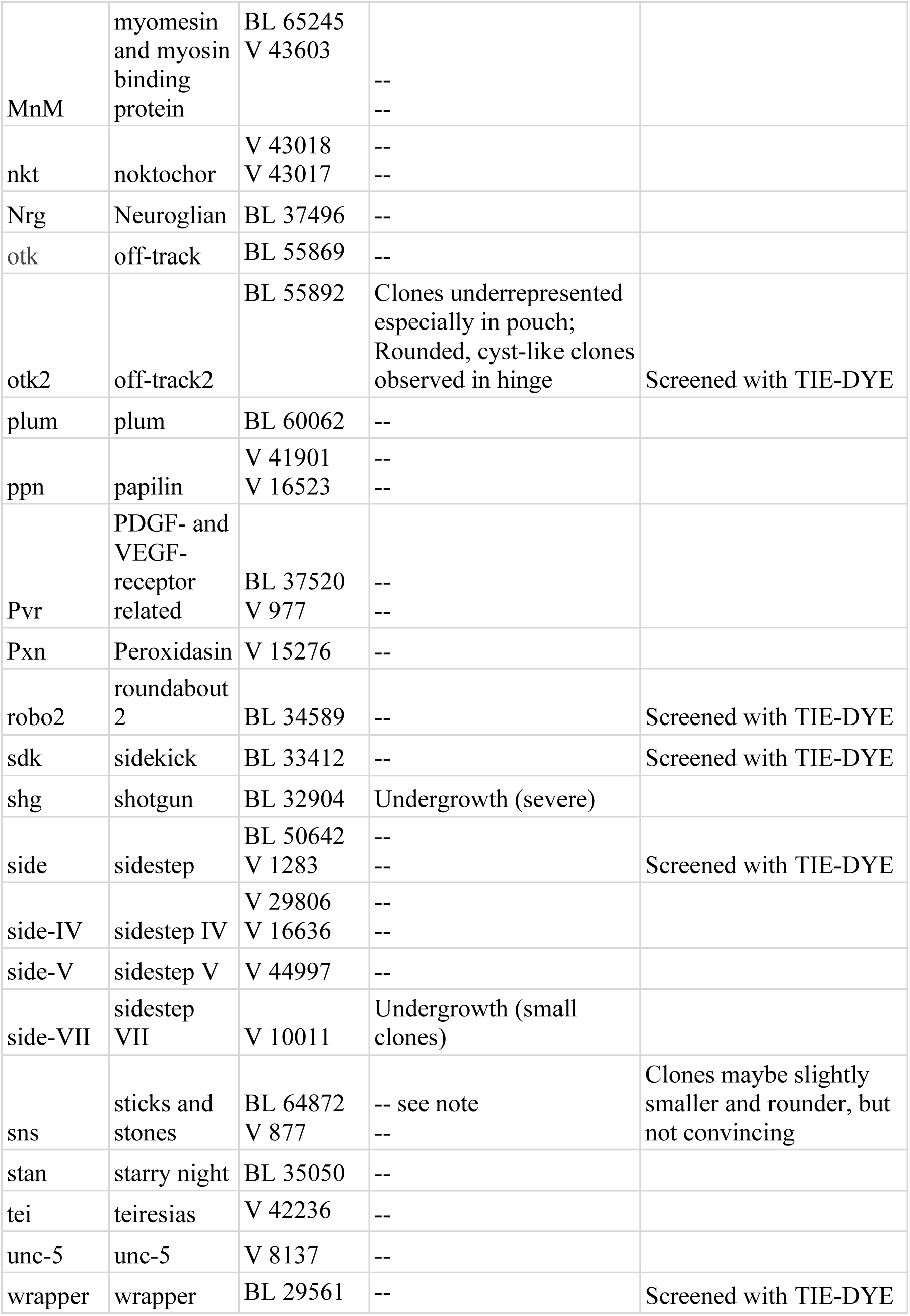
Results from lines which were screened.

